# *Cis*-regulation and natural selection in the evolution of human DNA methylation

**DOI:** 10.64898/2026.01.20.700710

**Authors:** Zhenzhen Ma, Alexander L. Starr, David Gokhman, Hunter B. Fraser

## Abstract

The vast collection of human-specific traits– such as our unique morphology, cognition, behavior, and diseases– has long been attributed to gene expression divergence between us and our closest living relatives, chimpanzees. Theory suggests that changes to *cis*-regulatory elements such as promoters and enhancers may drive evolutionary adaptation, and DNA methylation is a key factor in transcriptional *cis*-regulation. However, we still lack an understanding of 1) how species-specific methylation patterns arise; 2) their downstream effects; and 3) whether they are a common target of natural selection. In this study, we investigated these three questions. By combining a novel hypothesis testing framework with DNA methylation data from six human and chimpanzee cell types, as well as fused interspecies hybrid cells, we disentangled *cis*- vs. *trans*-acting methylation divergence across the genome. Across cell types, we found that methylation divergence is primarily driven in *cis*, which can be linked in some cases to nearby sequence variants such as CpG gains and losses. Although less common, regions with *trans*-acting methylation divergence were enriched for specific transcription factor (TF) binding motifs, suggesting a role of TFs such as FOXM1 in these differences. Having established these causes of methylation divergence, we then examined the functional consequences of differential methylation. Although methylation lacks a consistent relationship with transcription, we observed that associations between methylation and gene expression are stronger for genes with *cis*-regulatory divergence. Moreover, we identified lineage-specific selection shaping promoter methylation at the level of entire pathways including those associated with human-specific traits such as speech, cognition, and susceptibility to infection with hepatitis C. Collectively, our findings provide a mechanistic framework suggesting that DNA methylation may occupy a key position, mediating the effects of both *cis*- and *trans*-acting factors on transcriptional networks, including those contributing to human-specific traits.

## Introduction

While gene expression divergence has long been considered the primary driver of human evolution, identifying the molecular mechanisms underlying uniquely human traits remains challenging, particularly when they involve epigenetic modifications that control cell type-specific expression patterns^1–4^. Theory suggests that cell type-specific *cis*-regulatory variants may fuel evolutionary adaptation through their precise effects on individual genes within specific cell types, avoiding potentially deleterious consequences of broader *trans*-acting changes that affect multiple targets simultaneously^5,6^. However, while recent advances have enabled quantification of human-specific divergence in gene expression and chromatin accessibility, the role of DNA methylation—a key regulator of gene expression— in coordinating these changes and driving human-specific traits has not been fully explored^6–10^. DNA methylation, occurring primarily at CpG dinucleotides, is generally associated with transcriptional repression and provides stable, long-term regulatory control that has been linked to species-specific phenotypes in mammals^11^. Comparative methylome studies between humans and chimpanzees have revealed widespread tissue-specific methylation divergence, and linked human-specific hypomethylation to neurodevelopmental evolution and disease susceptibility^7,12–22^.

Studies of allele-specific expression (ASE) and chromatin accessibility measurements in interspecies hybrids provide a powerful approach for studying regulatory evolution by eliminating confounding factors inherent to cross-species comparisons, including differences in cell type composition, environmental conditions, and developmental timing^23–32^. Analogously, allele-specific methylation (ASM) in human-chimpanzee hybrid cells enables the isolation of *cis*-regulatory effects while controlling for shared *trans*-acting factors. *Cis* regulation of methylation is caused by factors such as sequence variants affecting methylation on the same allele as the variants, typically acting locally. In contrast, *trans* regulation of methylation involves diffusible factors such as DNA methyltransferases, transcription factors, or chromatin modifiers that can influence methylation patterns across many genomic loci (Figure 2A). Studying ASM enables direct quantification of the relative contribution of local versus diffusible factors to species-specific methylation patterns.

While previous studies of primate methylation evolution in post-mortem tissues and blood have provided valuable foundational insights^7,12–16,18–22,33,34^, several important questions remain to be addressed. These include: (1) generating comprehensive genome-wide methylation maps across multiple species and cell types; (2) distinguishing between *cis*-versus *trans*-regulatory mechanisms underlying species-specific methylation differences; (3) identifying *cis*-acting genetic variants and *trans*-acting factors underlying these differences; (4) identifying lineage-specific natural selection on methylation; and (5) linking this selection to gene expression and candidate traits. Addressing these gaps would enhance our understanding of how genetic variants impact methylation and in turn shape adaptive traits.

Human-chimpanzee tetraploid hybrid induced pluripotent stem cells (iPSCs), generated through *in vitro* cell fusion, provide a controlled experimental system for quantifying *cis*- and *trans*-regulatory contributions to methylation divergence by hosting both species’ alleles in an identical *trans*-environment. By measuring ASM and ASE across multiple cell types, this system generates genome-wide maps of cell type-specific regulatory differences that may contribute to evolutionary adaptation. In this study, we generated bisulfite sequencing data from human, chimpanzee, and hybrid iPSCs, cranial neural crest cells, dopaminergic neurons, skeletal myocytes, and hepatocytes, as well as from primary human and chimpanzee dental pulp stem cells. We combined these data with previously published^26^ and newly generated RNA-seq data from all six cell types.

Using these data, we investigated the causes of methylation divergence and quantified contributions of *cis*- and *trans*-acting mechanisms that establish interspecies differences in methylation patterns. We developed a hypothesis testing framework tailored to count-based proportional methylation data and extended a changepoint detection algorithm to identify differentially methylated regions (DMRs) under consistent *cis*- or *trans*-regulation. We show that *cis*-acting mechanisms predominantly drive genome-wide methylation divergence, with *cis*-regulated DMRs likely affected by nearby species-specific single-nucleotide variants. In contrast, *trans*-regulated DMRs were enriched for transcription factor binding motifs, including pioneer factors and chromatin remodelers that may mediate *trans*-acting divergence. Concordance between ASM and ASE across cell types revealed species-specific and context-dependent regulatory relationships linking methylation to gene expression. Finally, coordinated methylation-expression divergence in gene sets related to speech, cognition, and viral infection susceptibility connected lineage-specific selection to candidate phenotypes. In sum, our work provides a framework for understanding how divergent *cis*- and *trans*-acting factors converge on methylation patterns to produce species-specific traits.

## Results

### Measurement of DNA methylation across diverse cell types

To establish comprehensive methylation landscapes across evolutionarily relevant cell types, we differentiated iPSCs from two humans, two chimpanzees, and two fused tetraploid hybrids into four cell types (see Methods)^24,26^. We included an additional cell type, dental pulp stem cells, which are primary cells isolated from two human and two chimpanzee donors^24^. We then generated whole-genome bisulfite sequencing (WGBS) data from human and chimpanzee parental and hybrid cranial neural crest cells (CNCC) and dopaminergic neurons, as well as parental dental pulp stem cells (DPSC), complemented by reduced representation bisulfite sequencing (RRBS) data from induced pluripotent stem cells (iPSCs), skeletal myocytes (SKM) and hepatocytes (HEP) (Figure 1A). For cross-cell-type analyses combining WGBS and RRBS data, we restricted to CpG sites covered in all samples after phasing, totaling 561,762 shared CpG sites. For hybrid samples, reads were assigned to the human or chimpanzee allele using species-specific SNV positions; on average, 51% ± 0.4% of all aligned reads were allele-resolved across hybrid WGBS samples (DA and CNCC), and 10.6% ± 0.2% of aligned reads were allele-resolved across hybrid RRBS samples (IPSC, SKM and HEP). The lower rate in RRBS reflects its short read lengths and its enrichment for CpG-dense promoter regions, which are evolutionarily conserved and therefore depleted of informative human–chimpanzee SNVs (see "RNA-sequencing and allele-specific expression analysis" in Methods section). Principal component analysis and hierarchical clustering of methylation patterns showed that samples clustered primarily by cell type, then by species, and finally by parental (human and chimpanzee) versus hybrid system (Figure 1B, 1C and Figure 1—figure supplement 1). CNCC samples displayed a unique clustering pattern where parental vs. hybrid samples clustered separately, though examination of the dendrogram indicated this was due to a single human parental sample being a slight outlier (Figure 1B). The transcriptional distinctiveness of DPSCs relative to iPSC-derived cell types likely reflects their status as primary cells, which retain the epigenetic and transcriptional signatures of their tissue of origin and have not undergone reprogramming (Figure 1C). Known cell type-specific gene expression markers validated the correct identity of each differentiated cell type (Figure 1D and Figure 1—figure supplement 2)^35–49^. Overall, these results are consistent with our expectation that cell type is the predominant determinant of DNA methylation patterns, while species explains a much smaller proportion^7^. In other words, a human hepatocyte is much more similar to a chimpanzee hepatocyte than it is to a human dopaminergic neuron.

**Figure 1.**
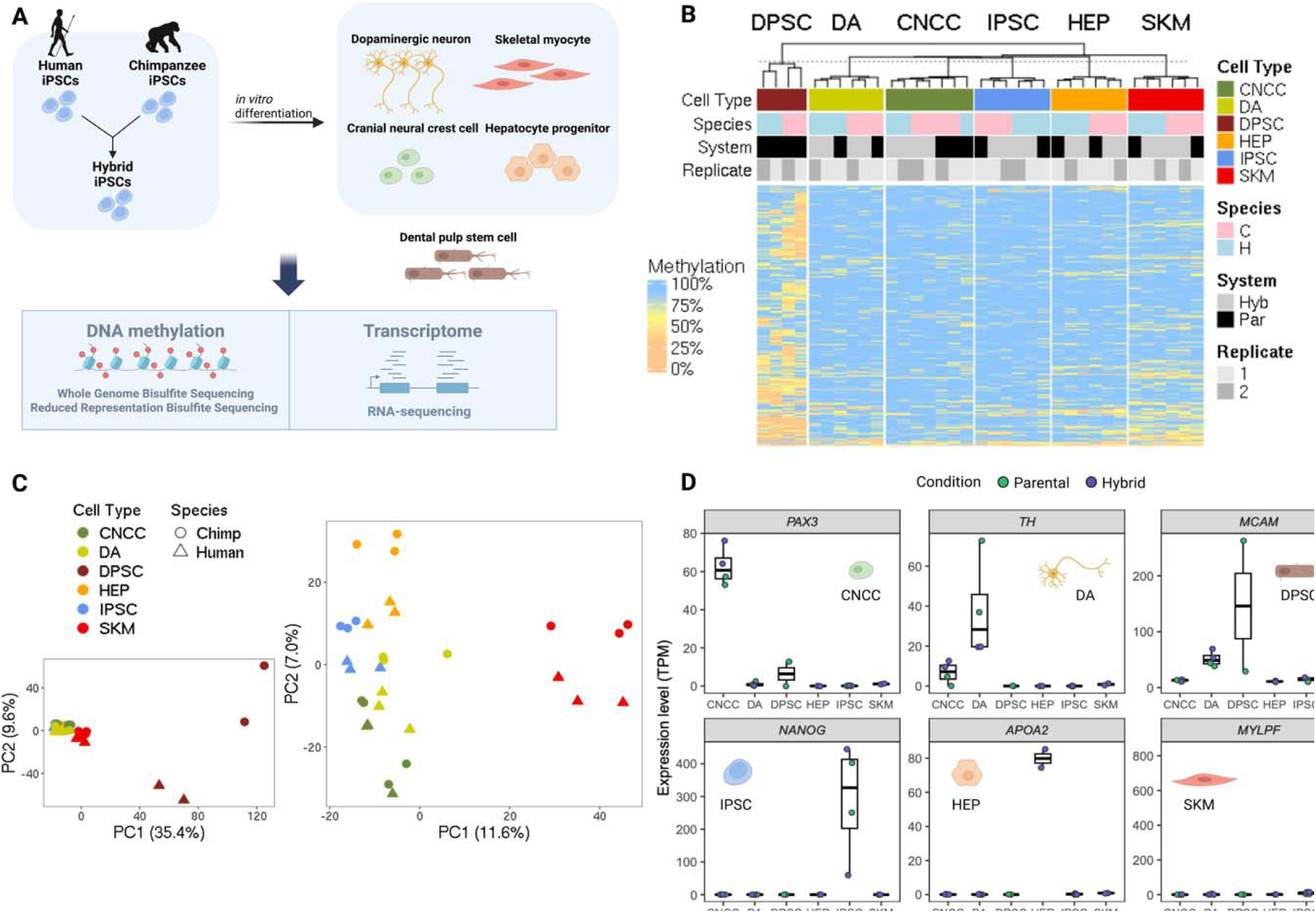
DNA methylation profiles across diverse human, chimpanzee, and hybrid cell types. (A) Four cell types were differentiated from human, chimpanzee, and hybrid induced pluripotent stem cells. The cell types span diverse body systems including skeletal myocytes for the musculoskeletal system, dopaminergic neurons for the central nervous system, cranial neural crest cells for craniofacial development and hepatocyte progenitors for the liver. We collected DNA methylation and RNA-seq data from these cell types from humans, chimpanzees and their hybrids. Dental pulp stem cells were also collected from human and chimpanzee adult tissues. (B) Clustering analysis and (C) PCA between samples for all cell types (left) and for all iPSC-derived cell types (i.e. excluding DPSC, right). Parent/hybrid stratifications are shown separately in Figure 1—figure supplement 1. Each replicate represents an independent biological differentiation from the indicated iPSC line. (D) Cell type specific markers of gene expression. Figure 1A was created using BioRender, and is published under a CC BY-NC-ND 4.0 license.

### Contribution of *cis* and *trans* regulation to DNA methylation divergence between human and chimpanzee

To understand the molecular mechanisms underlying species-specific methylation patterns, we first established a framework for distinguishing between *cis*- and *trans*-regulatory control of DNA methylation. By comparing methylation levels between human and chimpanzee alleles within the same cellular environment (hybrid cells), we can isolate *cis* effects from *trans* effects and determine which mechanism predominates in establishing species-specific methylation patterns (Figure 2A).

**Figure 2.**
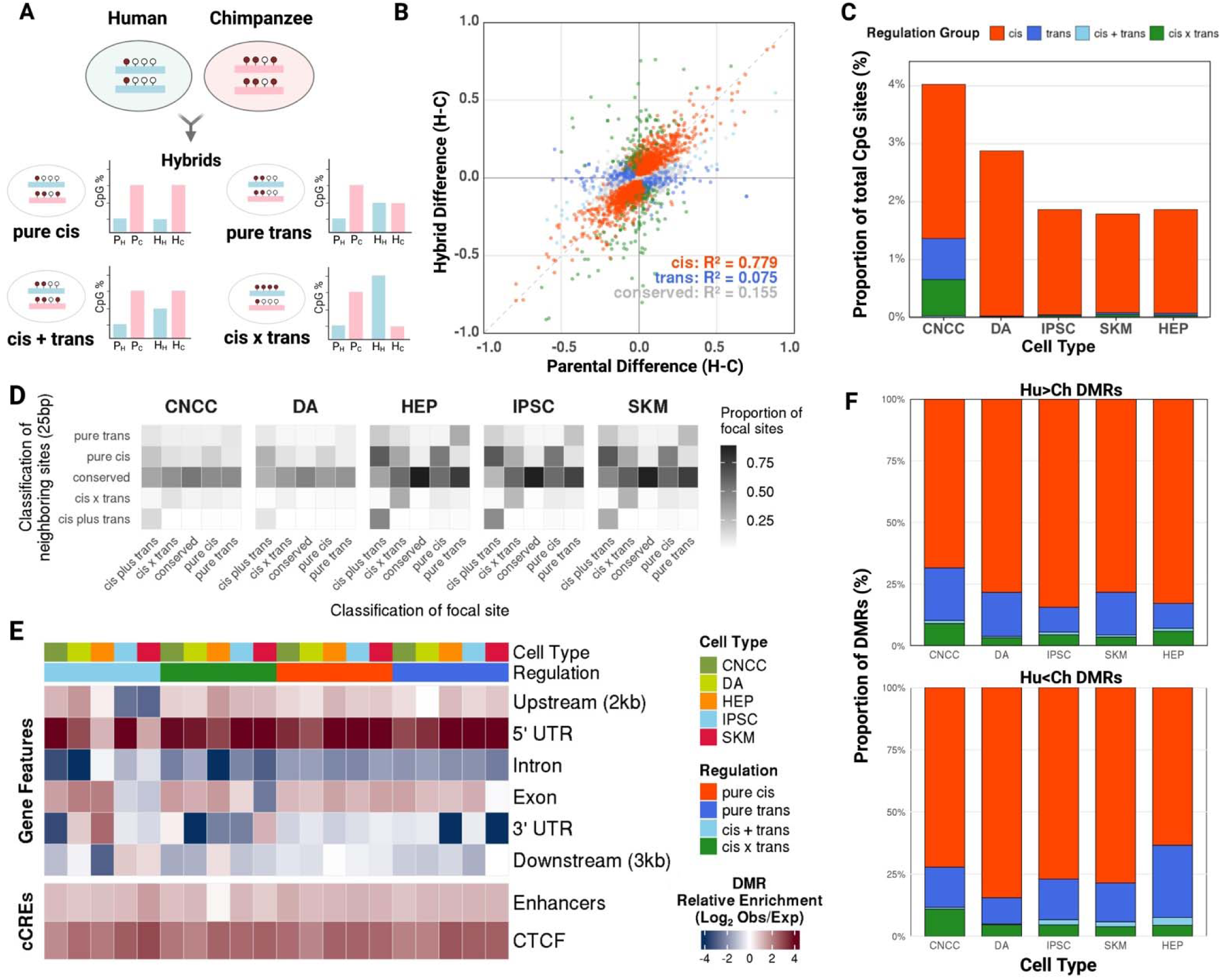
Contribution of *cis* and *trans* regulation to DNA methylation divergence between human and chimpanzee. (A) Interspecies differences in methylation levels per CpG site is compared between parental and hybrid systems and assigned a regulation category based on hypothesis testing. (B) Parental and hybrid methylation ratios in cranial neural crest cell promoters and the respective regulation category at FDR < 0.05 (color key in panel C). Parental and hybrid methylation differences are plotted as the per-sample average difference of fractional methylation. (C) Quantification of relative contribution of non-conserved regulation groups to whole-genome CpG sites across cell types at FDR < 0.05. (D) Heterogeneity of the distribution of CpG sites of all regulation groups. (E) Quantification of CpG regulatory clusters by overlap with genomic features of transcripts genome-wide as well as *cis*-regulatory elements including promoters, enhancers and CTCF-bound *cis*-regulatory regions (CTCF). (F) Quantification of relative contribution of non-conserved regulation groups to regulatory clusters across cell types (color key in panel C). Figure 2A was created using BioRender, and is published under a CC BY-NC-ND 4.0 license.

To quantify the relative contributions of *cis*- versus *trans*-acting factors in establishing human-chimpanzee methylation differences, we applied a beta-binomial model framework adapted from Hallgrímsdóttir et al. (2024)^50^ to classify differentially methylated regions based on allele-specific patterns in hybrid versus parental samples. To determine whether a CpG site reflects a difference in methylation due to *cis* or *trans* regulation, two hypothesis tests are used: one to evaluate the null hypothesis that the difference in methylation is due solely to *trans*, and the other solely to *cis* (see Methods). Each site is assigned to one of five mutually exclusive regulatory categories. *Cis*-only sites show allelic differences in hybrids that are also preserved in the parental comparison. *Trans*-only sites show allelic differences in parentals that are absent in hybrids. *Cis* + *trans* (additive) sites show allelic differences in both systems, with the same species methylated higher in both, but with the magnitude of interspecies difference reduced in hybrids, indicating a partial *trans* contribution acting in the same direction as *cis*. *Cis* × *trans* sites show allelic differences in both systems but with opposite directionality (one species higher in parentals, the other higher in hybrids). Conserved sites show no significant allelic differences in either system (Figure 2A and 2B). We investigated genome-wide methylation divergence by fitting the model to methylation levels at each CpG site (across 49,049,052 unique CpG positions covered by any library, 24,935,901 CpG sites shared across hybrid and parental samples across all cell types; see Supplemental File 1 for detailed statistics). In all cell types examined, most CpG sites (83–93%) had conserved patterns of methylation. Among divergent sites, nominally-classified cis-regulation was over twice as common as trans-regulation (Figure 2—figure supplement 1, Supplemental File 1). At FDR < 0.05, cis-regulation accounted for a mean of 91% of divergent sites and trans-regulation a mean of 4.5% across cell types, and cis remained the dominant category at every FDR cutoff tested (FDR < 0.25–0.01; Figure 2C, Figure 2—figure supplement 1–2, Supplemental Files 1–2). Summed across cell types, 663,368 CpG sites were classified as cis-regulated and 140,017 as trans-regulated at FDR < 0.05, out of 932,573 divergent sites among 24,935,901 tested sites (per-cell-type counts in Supplemental File 1). The genome-wide predominance of cis-regulation was further supported at the regional level (Figure 2B and Figure 2—figure supplement 1 and 3). This genome-wide predominance of cis-regulation indicates that local sequence differences between human and chimpanzee genomes are the primary drivers of methylation divergence, consistent with previous hypotheses of gene regulation that *cis*-regulatory changes may be less disruptive than broad *trans*-acting modifications^1–4,6^.

Because counts of classified sites weight all sites equally regardless of effect size and depend on statistical power (which is lower for trans effects), we additionally estimated the proportion of variance in interspecies methylation divergence explained by cis and trans components, analogous to heritability partitioning in population studies (see Methods). To reduce sampling noise, we pooled read counts across CpGs within 1-kb genomic bins (40,094– 2,025,021 bins per cell type). In CNCC, the only cell type with two parental replicates per species, cis effects explained 93% (95% CI 92–95%) of the variance in methylation divergence and trans effects 71% (70–73%). Because the two components were negatively correlated (covariance term −0.64; 95% CI −0.68 to −0.62), their individual contributions sum to more than 100%. In the remaining cell types, which have one parental line per species, we report bounds whose validity we confirmed in CNCC (see Methods). Cis effects explained 58–75% (DA), 65– 78% (HEP), 71–98% (IPSC) and 67–84% (SKM) of the variance in divergence, compared with 32–48%, 28–40%, 3–30% and 22–38% for trans effects, and covariance terms were small (−0.01 to −0.07). Cis effects therefore exceeded trans effects in every cell type under both bounds, and these results were consistent at a stricter coverage threshold and at single-CpG resolution (Supplemental File 1). The variance-based contribution of trans effects was nevertheless larger than suggested by counts of classified sites, which may reflect trans effects that are individually too small to be detected at single CpG sites but collectively contribute to divergence.

The regional distribution of sites from different regulation groups reveals a consistent pattern across cell types: conserved CpG sites are most likely to be near other conserved sites, while divergent CpG sites are likely to be near conserved sites as well as sites within the same regulation group (Figure 2D). This suggests that CpG sites within the same regulation group tend to form coherent clusters in the genome, and divergent CpG clusters typically have conserved sites scattered within. When conducting regional analyses, methylation data is usually pooled from multiple sites within a pre-defined genomic region^6–10^. However, this regional heterogeneity can affect classification as well as estimates of how much each regulatory mechanism contributes to divergence between species (Figure 2—figure supplement 2-3, Supplemental File 2, see Methods). Therefore, pooling data across regions should be approached carefully, as subsets of CpG sites within the same regulatory element can be controlled by distinct mechanisms. These findings underscore the importance of studying methylation divergence at high resolution.

To identify differentially methylated regions (DMRs) exhibiting consistent inter-species methylation directionality and minimal classification heterogeneity of regulation groups, we implemented a changepoint detection method with a tolerance threshold for regulatory homogeneity (see Methods). This method identifies genomic loci where co-localized CpG sites demonstrate uniform regulatory behavior punctuated by a pre-defined fraction of heterogeneous sites: non-conserved classifications showing consistent directional methylation bias, and conserved sites maintaining equal methylation levels between species. This method prioritizes DMRs that are controlled by the same mechanism and display coordinated methylation patterns (Supplemental File 3), thereby enriching for loci with interpretable regulatory mechanisms.

Notably, DMRs identified through this method are enriched in 5’ UTRs (just downstream of each transcription start site [TSS]) and CTCF-bound regions, supporting their functional relevance in gene regulation (Figure 2E). In total, we identified 9159 cis-DMRs and 2046 trans-DMRs across all cell types (see Methods and Supplemental File 3). The quantification of *cis*- vs. *trans*-DMRs yielded the same conclusion that *cis*-regulation outweighs *trans*-regulation in DNA methylation divergence (Figure 2F).

### The causes of *cis*- and *trans*-acting divergence in DNA methylation

To explore potential molecular mechanisms driving methylation divergence, we focused on two of many possible contributors: transcription factors (TFs) associated with trans-DMRs and sequence variants associated with cis-DMRs. TFs represent a major class of *trans*-acting factors capable of establishing and maintaining methylation patterns across the genome, often by recruiting methyltransferase or demethylase enzymes. Interspecies variation in TF expression levels, binding activity, or target site accessibility can propagate to downstream methylation differences at their binding sites. To identify TFs potentially driving species-specific methylation divergence through such *trans*-acting mechanisms, we performed TF binding site motif enrichment analysis on *trans-*DMRs (Figure 3A, Supplemental File 4 and 5)^51^. This analysis identified several candidate factors, including a pioneer factor FOXA2 (Benjamini-Hochberg corrected FDR = 3.7 × 10⁻ in DA Hu<Ch *trans*-DMRs) and a chromatin modifier FOXM1 (FDR = 0.002, 4.8 × 10⁻ , and 0.021 in CNCC, DA, and HEP Hu<Ch *trans*-DMRs, respectively), both known to actively reshape DNA methylation landscapes^52–58^.

**Figure 3.**
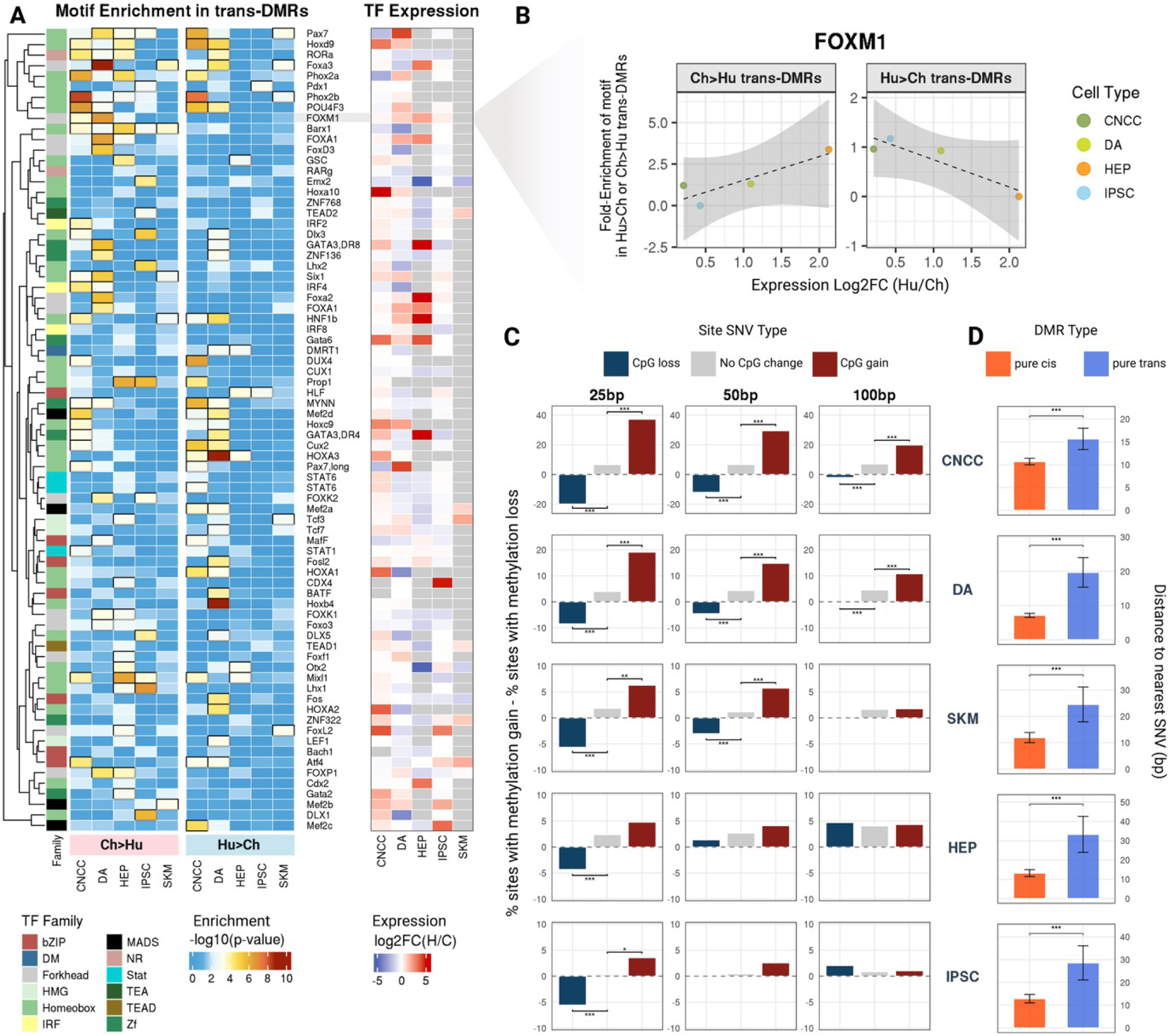
*Cis*- and *trans*- factors influencing species-specific methylation patterns. (A) HOMER motif enrichment of *trans-*DMRs in regions Hu>Ch and Ch>Hu methylated in humans (left panel), and differential expression of corresponding transcription factors in parents (right panel). (B) Fold-enrichments against genomic background of FOXM1, one of the top motifs enriched in Hu>Ch and Ch>Hu *trans-*DMRs, plotted against Hu and Ch relative gene expression levels in parents. (C) Three classes of SNVs—CpG gains in humans, CpG losses in humans, or no CpG change—and their effects on methylation of conserved neighboring CpG sites within ±25, 50, and 100 bp. The differential methylation patterns around CpG SNVs versus non-CpG-SNVs are assessed using two-proportion test (***: p < 0.001, **: p < 0.01, *: p < 0.05) (D) Distances of each *cis-*DMR vs *trans-*DMR to the nearest CpG SNV site (Mann-Whitney U test; ***: p < 0.001).

If species-specific levels of a TF are responsible for the enrichment of its predicted binding sites in *trans-*DMRs, then this enrichment may be strongest in cell types with greater species-specific differential expression of the TF. FOXM1 and FOXA2 each showed a pattern of this kind, with higher relative expression in one species accompanied by lower methylation at their predicted binding sites in that same species and cell type. FOXM1 was over 4-fold more highly expressed in human HEPs than in chimpanzee, and its predicted binding sites were highly enriched specifically for trans-DMRs with lower methylation in human HEPs; the three other cell types, in which human and chimpanzee expression of FOXM1 is more similar, showed weaker enrichment and little species-specific asymmetry in trans-DMR methylation directionality (Figure 3B). We observed a similar pattern for FOXA2 (Figure 3—figure supplement 1). We note that with only four cell types available per comparison, we are not adequately powered to test for a statistically significant relationship between differential TF expression and motif enrichment. We therefore present FOXM1 and FOXA2 as exploratory, hypothesis-generating examples rather than as evidence of a genome-wide pattern.

Other TFs showed different patterns of association between species-specific expression and *trans-*DMR methylation. For example, FOXP1 and OTX2 were most differentially expressed in HEPs, and their predicted binding sites were also most enriched in trans-DMRs in this same cell type (Figure 3—figure supplement 1). However, unlike FOXM1 and FOXA2, binding sites for FOXP1 and OTX2 showed similar patterns of enrichment across cell types for both Hu>Ch and Ch>Hu methylated *trans-*DMRs. This lack of directionality may suggest that these transcription factors are not restricted to either demethylation or methylation, perhaps having a more indirect or context-specific role in regulating the epigenetic landscape^59–61^, though further work is required to test this hypothesis. Altogether, our *trans-* DMR analysis nominated many TFs that may contribute to methylation divergence, including several with *trans-*DMR binding site motif enrichment associated with the TF’s differential expression that we highlight for future experimental testing.

To identify the molecular mechanisms driving *cis*-regulatory methylation differences, we then examined whether local sequence variants could explain *cis*-acting methylation divergence. CpG dinucleotides are the primary targets of DNA methylation in mammalian genomes, and their presence, density, and spacing are known to influence local methylation patterns^62–64^. Therefore, single nucleotide variants (SNVs) that either create or disrupt CpG dinucleotides represent a plausible mechanism by which sequence divergence could drive species-specific methylation differences. To investigate this hypothesis, we systematically analyzed SNVs and their associated methylation differences in neighboring conserved CpGs within windows of 25, 50, and 100 base pairs upstream and downstream of each SNV. Our analysis revealed that SNVs affecting CpG dinucleotides—but not other SNVs—are associated with altered methylation patterns of nearby CpG sites up to 50 base pairs from the variant site (Figure 3C). Notably, the effect diminished with increasing distance from the SNV, with the strongest methylation changes observed within 25 base pairs and progressively weaker effects at 50 base pairs. Beyond 50 base pairs, the magnitude of influence of CpG SNVs on neighboring methylation patterns became negligible, suggesting a limited spatial range for these local sequence effects. This distance-dependent relationship indicates that sequence variants affecting CpG density and spacing might influence the local recruitment or activity of DNA methyltransferases or demethylases. We note that CpG SNVs represent just one of many possible mechanisms for cis-methylation divergence; other classes of variants, such as those disrupting transcription factor binding sites, may also contribute and warrant future investigation.

This model predicts that while *cis-*DMRs should often be close to CpG SNVs, *trans-* DMRs—which arise from regulatory mechanisms independent of nearby sequence variation— should not be. To test this, we compared the genomic distance between each DMR and its nearest CpG SNV. Consistent with our model, we found consistently shorter distances for *cis-* DMRs than *trans-*DMRs across all cell types examined: Median distances to the nearest CpG SNV were 24 ± 6 bp for trans-DMRs vs. 11 ± 2 bp for cis-DMRs (Figure 3D; Mann-Whitney p < 0.001 for all comparisons)^65^. This difference suggests that *cis-*DMRs tend to occur near CpG sequence variants and are likely directly influenced by local changes in CpG content, while trans-DMRs show no preferential association with CpG variants, as expected given that trans-DMRs are by definition not driven by local sequence variation.

To further test the effects of CpG SNVs on local methylation, we assessed whether the same SNVs show consistent methylation effects across multiple cell types. CpG SNVs associated with a cis-DMR in one cell type were significantly more likely to be associated with a cis-DMR in other cell types than expected by chance, consistent with a shared mechanistic basis for cis-methylation divergence across cell types (Figure 3—figure supplement 2). This genomic organization is consistent with *trans-*DMRs being regulated by diffusible factors such as transcription factors rather than by local sequence variants, supporting a model in which *cis*- and *trans*-acting mechanisms operate through fundamentally different molecular pathways to generate species-specific methylation patterns.

### Cell type- and species-specific gene expression as a consequence of methylation divergence

Given the well-established repressive effect of promoter methylation on gene expression, we assessed whether methylation differences contribute to expression divergence by analyzing genome-wide correlations between allele-specific promoter methylation and expression levels across different genomic contexts (Figure 4A)^66–68^. We first examined whether *cis*-acting variants drive promoter methylation consistently across cell types or in a context-dependent manner. We found that while some promoters are allele-specifically methylated across all cell types, many more showed cell type-specific *cis*-methylation patterns, suggesting that promoter methylation regulation is often context-dependent (Figure 4B and 4C). However, while DMRs consistently detected across cell types provide strong evidence of shared regulation, DMRs detected in only one cell type may reflect either genuine cell type-specificity or insufficient power to detect them elsewhere. Our estimate of cell type-specific regulation therefore represents an upper bound on the degree of cell type-specific methylation regulation.

**Figure 4.**
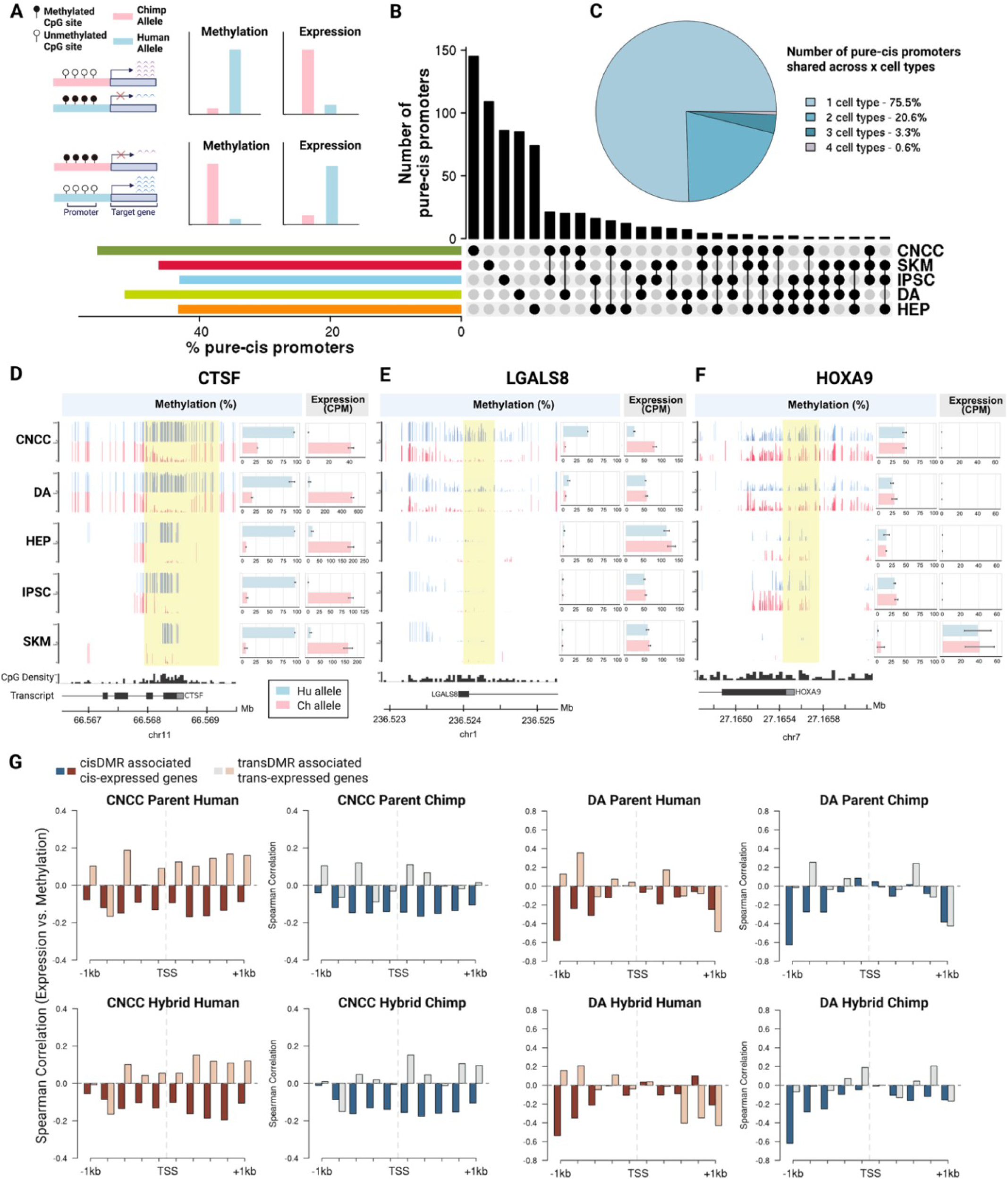
Human-chimpanzee allele-specific methylation can be associated with allele-specific expression. (A) Illustration of the transcriptional consequences of DNA methylation explored in this study. DNA methylation, as a stable *cis*-acting regulatory mark, often represses transcription. In a hybrid, allele-specific methylation can lead to allele-specific gene expression, although both methylation and its effects on transcription can be context-dependent. (B) We identified many promoters with allele-specific methylation in each cell type, most of which showed allele-specific methylation in only one cell type. (C) Quantification of promoters with pure-*cis* regulation of methylation across cell types. (D, E, F) Examples of gene expression-methylation correspondence. Methylation tracks show fractional methylation (mC/coverage × 100%) of individual CpGs. Expression values are in TMM (Trimmed Mean of M-values)-normalized CPM (counts per million). (D) *CTSF* as an example of species-specific, cell type-agnostic DMR, (E) *LGALS8* as an example of cell type-specific allele-specific methylation, and (F) *HOXA9* as an example of cell type-specific but species-agnostic methylation patterns. (G) Genome-wide profile of expression-methylation correlations (binned by distance of methylated region to the TSS) for genes with both promoter methylation and expression under *cis*-regulation (dark red and dark blue) or *trans*-regulation (light brown and light gray). Detailed statistics are provided in Supplemental File 6. Figure 4A was created using BioRender, and is published under a CC BY-NC-ND 4.0 license.

Several specific examples illustrate different patterns of methylation-expression coordination (Supplemental Files 5 and 6; see Methods). The promoter of *CTSF* is a cell-type agnostic *cis-*DMR with consistent Hu>Ch methylation across cell types corresponding to coordinated Ch>Hu expression (Figure 4D and Figure 4—figure supplement 1). In contrast, a putative enhancer in *LGALS8* represents a case where methylation divergence is confined to specific cellular contexts. The candidate enhancer overlaps with a *cis-*DMR with Hu>Ch methylation and *LGALS8* exhibits allele-specific Ch>Hu expression exclusively in CNCCs, while having no divergence in methylation or expression in other cell types (Figure 4E and Figure 4—figure supplement 2). A third type of pattern is exemplified by the *HOXA9* promoter, with cell type-specific hypomethylation corresponding to cell type-specific expression only in SKM (Figure 4F). These examples illustrate how different regulatory mechanisms can produce distinct patterns of methylation-expression coordination and how epigenetic changes may contribute to both species-specific and cell type-specific regulatory evolution.

Previous studies have reported weak but significant correlations between DNA methylation and gene expression divergence in primates^7,12,14,16,69^. Consistent with these findings, our genome-wide estimates of human–chimpanzee transcriptome divergence produced similar weak but significant correlations with methylation divergence (Figure 4—figure supplement 3 and Supplemental File 6). Our comparison of CpG island (CGI) vs. non-CGI promoters also yielded trends consistent with past studies showing that CGI promoter methylation correlates more strongly with gene expression repression than does non-CGI promoter methylation^70,71^ (Figure 4—figure supplement 4).

However, such genome-wide correlations may mask stronger relationships that occur within specific regulatory contexts where methylation and expression are more directly coupled. We hypothesized that our hybrid system could help identify genes exhibiting stronger methylation–expression coupling by stratifying them based on regulatory mechanisms. We reasoned that genes whose methylation and expression are both influenced by *cis*-acting variants— potentially driven by the same underlying genetic differences— should display tighter coupling than genes regulated through independent mechanisms.

To test this, we evaluated the extent to which methylation and expression show a negative relationship (i.e., higher methylation associated with lower expression) consistent with a repressive effect of methylation. In both CNCC and DA hybrid cells, we found that genes with *cis*-regulated expression were more likely to exhibit repressive patterns than those with *trans*-regulated expression (Figure 4—figure supplement 5). This is consistent with DNA methylation acting as an upstream *cis*-regulatory mechanism: genes with stronger *cis*-regulation are expected to show tighter methylation-expression coupling due to the greater contribution of local regulatory factors, including methylation itself. Further restricting the analysis to genes with concordant regulatory mechanisms revealed that genes with both methylation and expression being *cis*-regulated (“*cis*–*cis*” genes; Supplemental File 6) exhibited significantly stronger negative methylation–expression correlations than “*trans*–*trans*” genes (Figure 4G, see Supplemental File 6 for detailed statistics). This supports the hypothesis that methylation– expression coupling is strongest when both are driven by local genetic variation, perhaps through shared causal variants. Different shapes of the methylation–expression correlation profiles around the TSS in each cell type may reflect differences in the chromatin landscape and transcription factor occupancy that modulate the functional impact of promoter methylation in each cell type context; a deeper mechanistic explanation is left for future work.

### The sign test identifies lineage-specific selection on DNA methylation between human and chimpanzee

To identify pathways that have evolved under lineage-specific selection, we used a two-step binomial sign test approach that tests for non-random directional bias in methylation-expression patterns in hybrids^1,23,24,26,72–76^. The use of hybrid cell lines is critical for this analysis as it ensures that different promoters represent independent observations, controlling for *trans*-acting factors and environmental variation that could otherwise confound the detection of *cis*-regulatory effects. This test operates on the logic that neutral evolution should act like flipping a fair coin: if human-biased and chimpanzee-biased changes occur at equal background frequencies, then methylation and expression changes of genes within a pathway would accumulate randomly, showing no preference for human-biased versus chimpanzee-biased directionality. In contrast, lineage-specific selection would bias the coin flip, potentially causing functionally related gene sets to consistently favor one direction over the other rather than splitting evenly by chance.

To maximize our ability to find phenotypically impactful divergence, we adapted the standard sign test to account for directionality of both methylation and gene expression. In the first step, we applied a standard binomial sign test to all 6,283 promoters showing ASM, counting the number of genes showing human-biased versus chimpanzee-biased methylation changes within each gene set. The background probability for this test was defined as the overall fraction of promoter ASM in each cell type that was human-biased. This initial analysis identified gene sets showing significant enrichment for directional methylation changes, establishing candidates for lineage-specific regulatory divergence. In the second step, we restricted our attention to genes showing a negative association between ASM and ASE. This subset represents cases where methylation changes are most likely to have a simple and direct repressive effect on transcription, filtering out more complex regulatory scenarios where methylation and expression changes may be uncoupled or subject to competing regulatory influences.

Gene sets showing significant deviation from neutral expectation in the ASM/ASE concordance test (nominal binomial p-value < 0.05 and FDR < 0.25 after multiple test correction; see Methods), with supporting evidence of directional bias in the methylation-only sign test (nominal binomial p-value < 0.05), are interpreted as likely targets of lineage-specific selection on promoter methylation that could directly impact gene expression (Figure 5, Supplemental File 7). However, as indicated by the FDR, we do expect some fraction of these candidate gene sets to be false positives. Compared to a standard sign test that focuses solely on directionally biased gene expression^72^, this more stringent two-step filtering approach may identify higher-confidence candidates where regulatory divergence is reflected at both the methylation and expression levels, establishing a potential mechanistic connection to phenotype.

**Figure 5.**
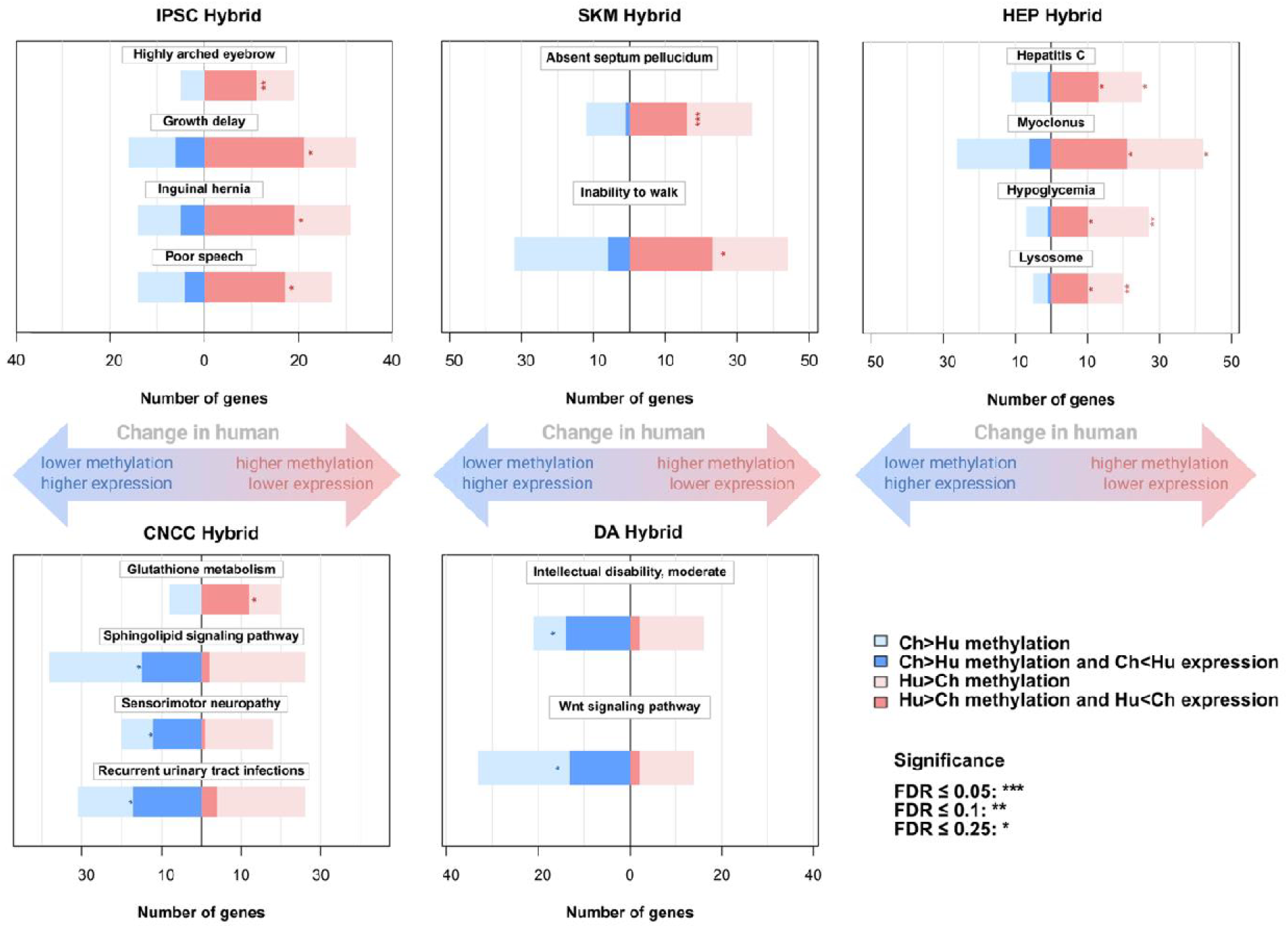
Gene sets with evidence of lineage-specific selection on methylation and gene expression. The length of the bars indicates the number of genes in each gene set with allele-specific expression or methylation in each direction (human or chimpanzee-biased). The bars in darker colors represent the number of genes where higher methylation is associated with repressed gene expression, whereas lighter colors represent genes where higher methylation is associated with increased gene expression.

Among gene sets from Kyoto Encyclopedia of Genes and Genomes (KEGG) and Human Phenotype Ontology (HPO)^77,78^, we identified several with clear relevance to human-specific traits. In iPSC hybrids, multiple gene sets including "poor speech," "highly arched eyebrow," and "growth delay” all showed Hu>Ch methylation and Ch>Hu expression (Figure 5, Figure 6A and Figure 6B). In the SKM hybrids, genes related to “inability to walk” showed significant Hu>Ch methylation and Hu<Ch expression (Figure 5). In DA hybrids, genes involved in “intellectual disability, moderate” show significant Hu<CH methylation and Hu>Ch expression (Figure 5 and Figure 6C). In HEP hybrids, we identified the “Hepatitis C” gene set with significant Hu>Ch methylation and Hu<Ch expression (Figure 5 and Figure 6D). In primary DPSC samples, we identified significant bias toward Hu<CH methylation and Hu>Ch expression in genes related to tooth eruption timing, consistent with known differences in dental development between humans and chimpanzees, where human permanent molars emerge approximately 6 years apart compared to 3-4 years in chimpanzees (Figure 5—figure supplement 1)^79–81^. However, the DPSC sign test results should not be interpreted as evidence of lineage-specific selection due to the use of parental rather than hybrid cell lines, which may introduce dependencies among observations due to trans-acting effects, potentially inflating statistical significance. For all other cell types (iPSC, DA, SKM, HEP, CNCC), the sign test results are based on hybrid lines and can be interpreted as evidence of lineage-specific selection, as the hybrid system controls for trans-acting confounders.

**Figure 6.**
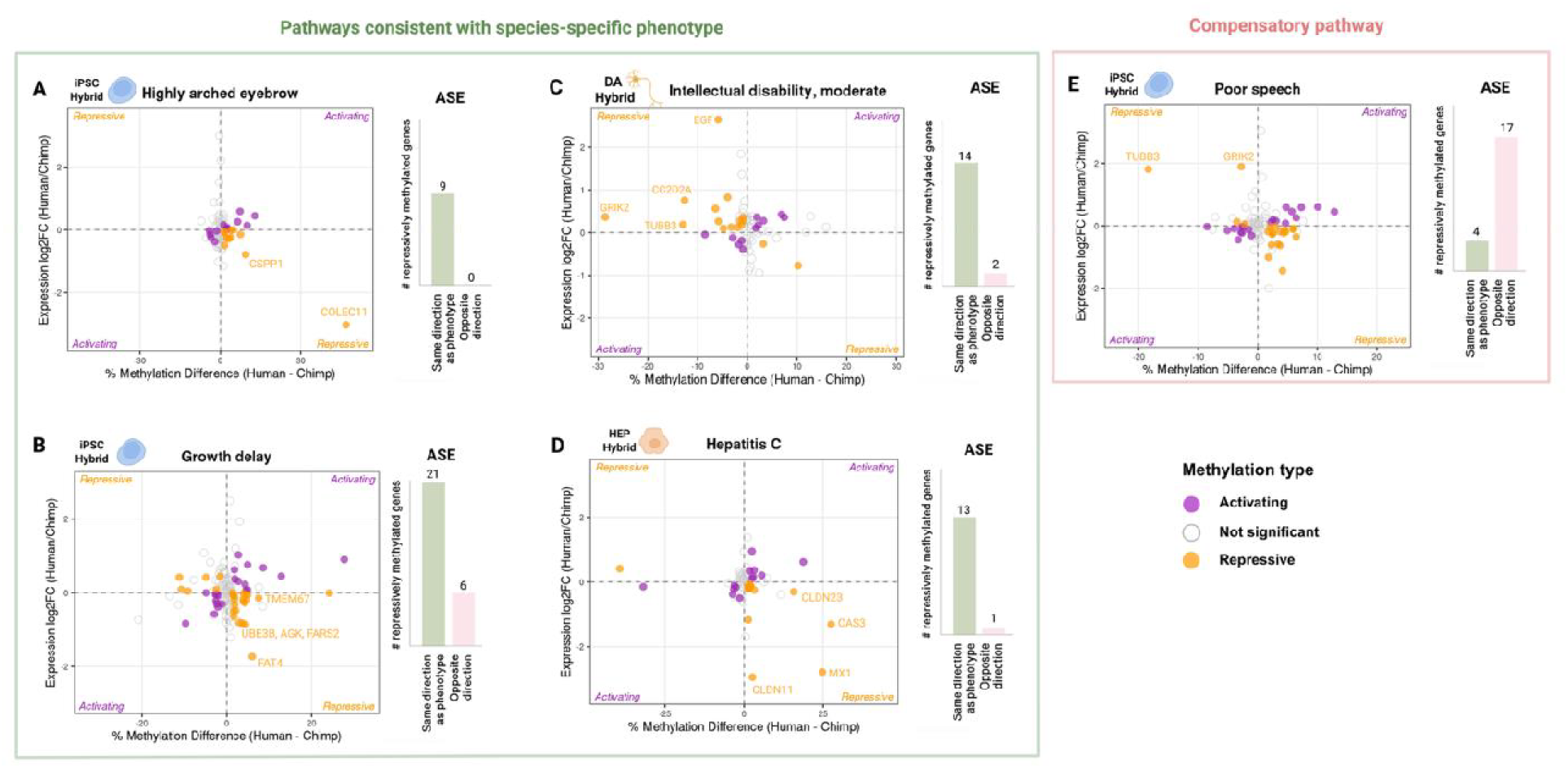
Directional bias in allele-specific methylation and gene expression identifies candidate genes for human-specific phenotypes. Allele-specific expression (ASE) is plotted against allele-specific methylation (ASM). Orange dots represent genes consistent with repressive methylation (e.g. Hu>Ch ASE and Hu<Ch ASM) and purple dots represent genes consistent with activating methylation (e.g. Hu>Ch ASE and Hu>Ch ASM). Bar plots represent counts of genes with repressive methylation that exhibit expression changes in directions either consistent with (light green) or opposite to (light pink) what would be expected based on phenotypic differences between humans and chimpanzees. A) The “highly arched eyebrow” gene set in iPSC hybrids shows consistent Hu<CH ASE and Hu>Ch ASM for genes with repressive methylation. B) The “growth delay” gene set in iPSC hybrids shows significant bias towards Hu<CH ASE and Hu>Ch ASM for genes with repressive methylation. C) The “intellectual disability, moderate” gene set in DA shows consistent Hu>Ch ASE and Hu<Ch ASM for genes with repressive methylation, with genes including *GRIK2, TUBB3, EGF and CC2D2A* being among the genes with highest magnitude ASE and ASM. D) The “Hepatitis C” gene set in HEP shows consistent Hu<CH ASE and Hu>Ch ASM for genes with repressive methylation. E) The “poor speech” gene set in iPSC hybrids represents a potentially compensatory pathway where most genes with repressive methylation show Hu<CH ASE and Hu>Ch ASM, whereas the genes with highest magnitude of ASE and ASM (*TUBB3* and *GRIK2*) show Hu>Ch ASE and Hu<Ch ASM.

We then analyzed individual genes within each candidate gene set to assess whether expression bias in associated phenotypes aligns with known human-chimpanzee trait differences, focusing on genes showing repressive methylation. Multiple gene sets demonstrated directional bias in gene expression consistent with the direction of phenotypic divergence found between humans and chimpanzees. For instance, in the "highly arched eyebrow" gene set—where most genes show human down-regulation in iPSC hybrids— *COLEC11* (which has 8-fold lower expression from the human allele) stands out as an outlier in magnitude. *COLEC11* acts as a guidance cue in directing neural crest cell migration during early development, and in humans its loss-of-function (LOF) causes craniofacial dysmorphism including highly arched eyebrows and underdeveloped brow ridge, both of which are traits unique to humans (Figure 5 and Figure 6A)^82^. In the "growth delay" pathway, most genes with repressive methylation (whose LOF in humans and other species leads to delayed growth) show lower expression from their human alleles, consistent with humans’ substantially delayed growth pattern compared to chimpanzees (Figure 6B)^83,84^. Many of the genes in DA whose LOF causes "intellectual disability, moderate" may have contributed to the evolution of human cognitive abilities. For example, the high magnitude genes with Hu>Ch ASE and Hu<Ch ASM such as *GRIK2*, *TUBB3, EGF* and *CC2D2A* are essential for excitatory synaptic plasticity and long range axon wiring (Figure 6C)^85–90^. In addition, most genes whose inhibition or LOF increases susceptibility or severity of Hepatitis C (e.g. *MX1* and *OAS3*) show lower expression from human alleles in HEP, consistent with human’s heightened vulnerability to Hepatitis C viral infections (Figure _6D__)_91–93.

Interestingly, we also discovered gene sets where the direction of most gene expression differences predicted the opposite of a known human/chimpanzee phenotypic difference. For example, in the “poor speech” gene set, where loss-of-function is associated with impaired speech in humans, we counterintuitively observed that most genes were down-regulated in humans compared to chimpanzees. However, the two genes with the largest fold-changes, *TUBB3* and *GRIK2*, showed higher expression from the human allele (Figure 6E). Notably, these two genes are also among the large-magnitude genes in the predominantly human up-regulated “intellectual disability, moderate” pathway, which does align with species-specific phenotype (Figure 6C). This suggests the speculative possibility of compensatory evolution in which large, pleiotropic effects of a few genes are counterbalanced by smaller changes in many genes (Figure 6C and Figure 6E; also see Discussion). Our previous study found that genes affecting voice box show particularly extensive down-regulation in humans, consistent with our observation in the “poor speech” pathway^24^.

Collectively, these results provide the first evidence for lineage-specific selection on DNA methylation. Our use of hybrid cell lines was critical to this discovery, as it allowed us to isolate *cis*-acting regulatory changes from confounding *trans*-acting and environmental effects. Isolating *cis*-acting changes is critical because the sign test requires that each promoter represents an independent observation; in hybrid cells, each allele’s methylation is determined by its own *cis*-regulatory sequence, making observations across promoters genuinely independent and thus satisfying the assumptions of the binomial test. This approach reveals that polygenic selection has acted on entire sets of functionally related genes whose expression is modulated through DNA methylation. Importantly, by connecting lineage-specific methylation divergence to expression changes within functionally coherent pathways, we can generate plausible hypotheses linking regulatory evolution to human-specific phenotypes. These include craniofacial morphology, dental development, viral infection susceptibility, and neurodevelopmental characteristics underlying speech and cognition (Figures 5 and 6, Figure 5—figure supplement 1).

## Discussion

Our investigation using human-chimpanzee hybrid tetraploid cells revealed that *cis*-regulatory mechanisms are the primary drivers of genome-wide methylation divergence between humans and chimpanzees across multiple cell types, including dopaminergic neurons, skeletal muscle, iPSCs, hepatocytes, and cranial neural crest cells. This finding aligns with previous studies highlighting the predominant role of *cis*-regulatory elements in evolutionary adaptations, as they enable precise, context-specific gene regulation while minimizing the potentially deleterious pleiotropic effects of *trans*-acting factors^6^. We demonstrated that most differentially methylated sites arise through *cis*-regulatory mechanisms, with *trans*-regulation and *cis*-*trans* interactions playing secondary roles. We identified clusters of CpG sites with consistent regulatory mechanisms, forming coherent differentially methylated regions (DMRs). These DMRs predominantly overlap with functionally relevant genomic features including 5’ UTRs and upstream promoter regions, underscoring their regulatory significance.

Our mechanistic analysis revealed distinct pathways through which *cis*- and *trans*-regulatory factors establish species-specific methylation patterns. For *cis*-regulation, we identified CpG-altering sequence variants as a direct mechanism linking genomic divergence to epigenetic differences. These variants influence methylation patterns of neighboring CpG sites with effects diminishing with genomic distance. Importantly, *cis-*DMR clusters are significantly closer to CpG SNVs than *trans-*DMRs across all cell types, providing evidence that local sequence variation directly shapes *cis*-regulatory methylation patterns, perhaps through altered CpG density affecting recruitment of DNA methyltransferases or demethylases.

Our analysis of allele-specifically methylated promoters revealed that the majority showed cell type-specific patterns. Our examples illustrate this diversity: the *CTSF* promoter exhibits consistent cross-cell-type *cis*-methylation divergence (Figure 4—figure supplement 1) with corresponding expression changes; an enhancer near *LGALS8* shows cell type-specific allele-specific methylation and overlaps with a homogeneous *cis-*DMR exclusively in cranial neural crest cells; and the *HOXA9* promoter displays cell type-specific methylation patterns with corresponding context-dependent expression effects. These findings highlight how the broader regulatory context of each cell type impacts the effects of *cis*-regulatory variants.

Our genome-wide correlation analyses confirmed the well-known pattern that while promoter methylation generally shows statistically significant inverse relationships with gene expression, these correlations are weak when examined across all genes. However, by leveraging our hybrid cell system to stratify genes based on regulatory mechanism, we uncovered significantly stronger negative correlations between methylation and expression when both are cis-regulated compared to genome-wide estimates or *trans*-regulated genes. This finding suggests that the repressive effects of species-specific methylation are strongest when both layers of regulation are driven in *cis*, perhaps because *cis*-regulated methylation changes tend to have larger effect sizes than *trans*-regulated changes.

Our two-step binomial sign test approach in hybrids identified multiple gene sets showing coordinated directional shifts in methylation and expression that significantly exceed neutral expectations. Many of the pathways we identified are related to known human-chimpanzee phenotypic differences: intellectual disability in hybrid DAs, growth delay and highly arched eyebrow in hybrid iPSCs, hepatitis C in hybrid HEPs, and tooth eruption timing in parental DPSCs (Figure 5, 6, Figure 5—figure supplement 1). Specifically, the evidence rejecting a neutral null model of *cis*-regulatory evolution consists of statistically significant directional bias in both promoter methylation and gene expression within functionally coherent gene sets, assessed using a two-step binomial sign test with permutation-based FDR correction. These findings represent the first evidence for coordinated directional divergence in DNA methylation and identify candidate genes under lineage-specific selection. By restricting analysis to genes showing negative methylation-expression associations—where methylation most likely exerts direct transcriptional repression—we identified high-confidence candidates where epigenetic divergence has likely functional consequences. Our results suggest that *cis*-regulatory changes in DNA methylation represent an important substrate for adaptive evolution, complementing protein-coding and other *cis*-regulatory changes in generating human-specific traits.

We developed several methodological innovations tailored to the unique challenges of comparative epigenomics. First, our beta-binomial framework accounts for the count-based nature of bisulfite sequencing data and explicitly models allele-specific patterns in hybrid versus parental contexts. Second, our changepoint detection algorithm identifies coherent DMRs with minimal regulatory heterogeneity, prioritizing functionally organized regulatory units over isolated CpG sites. Third, the two-step sign test we introduce provides a method to identify lineage-specific selection that links methylation and corresponding changes in gene expression. These methods can be readily applied to other taxa where hybrid organisms can be generated.

However, our study also has important limitations. First, while hybrid cells provide powerful controls for *trans*-acting variation, in vitro differentiation protocols do not fully recapitulate the complexity and temporal dynamics of in vivo development. Therefore, cell lines are limited in their ability to capture species-specific methylation or expression profiles that are specific to a certain developmental stage or in vivo context. Second, pathway analysis of DPSCs used primary human and chimpanzee samples rather than hybrids, with *trans*-acting factors potentially inflating statistical significance in the sign test; these results should therefore be interpreted not as evidence for positive selection, but rather as an interesting pattern that could have important connections to phenotypic differences between humans and chimps. Third, while we identified associations between transcription factor binding motifs and *trans-*DMRs, definitive demonstration of causality requires targeted experimental perturbations such as base editing (for CpG-disrupting SNVs) or TF knockdown/overexpression experiments (for trans-acting TFs). Fourth, our study examined a limited number of cell types, and methylation divergence in most tissues remains unexplored. Fifth, our study lacks an outgroup species; therefore, we cannot definitively attribute observed differences to the human lineage specifically, and our use of the term "human-specific" refers to differences between humans and chimpanzees rather than changes unique to the human lineage. Sixth, while we focused on CpG SNVs as a mechanism for cis-methylation divergence, other classes of variants—such as transcription factor binding site mutations—could also influence local methylation patterns and represent an important avenue for future investigation.

Comparative gene expression analyses between humans and closely related species offer a powerful way to explore the molecular basis of species-specific traits and disease susceptibilities. However, it remains challenging to infer phenotypic differences through the lens of human clinical ontologies. To start with, HPO terms are largely derived from pathological human data and predominantly capture phenotypic consequences of loss-of-function mutations in a clinical context. When comparing expression levels between species, one might assume that lower expression corresponds to a loss-of-function-like phenotype. However, the relationship between expression dosage and function is typically nonlinear^94^, and complete gene knockouts in a model organism or loss of function mutations in patients are not equivalent to moderate downregulation. This fact may help explain cases where our sign test results suggest the opposite directionality as compared to human/chimpanzee phenotypic differences (Figure 6E). Alternatively, humans and chimpanzees may achieve a similar phenotype through distinct configurations of regulatory networks. For example, an adaptive down-regulation of a particular gene may be compensated for by up-regulation of other genes in the same pathway, maintaining overall pathway function. In other words, compensatory changes may complicate the relationship between gene expression and phenotype, as may be the case for the “poor speech” gene set (Figure 6E)^23^.

Several important directions emerge from this work. First, extending this framework to additional cell types and developmental time points would reveal whether the predominance of *cis*-regulation we observe is universal or varies across developmental contexts. Longitudinal sampling during differentiation could identify critical windows when methylation patterns are established and reveal whether human-chimpanzee differences emerge gradually or through discrete developmental transitions. Second, integrating chromatin accessibility (ATAC-seq) and histone modification (ChIP-seq) data would provide a more complete picture of the regulatory landscape, clarifying how DNA methylation interacts with other chromatin features to establish gene expression patterns. Third, direct examination of transcription factor binding using techniques like CUT&RUN in both species could validate the predicted relationships between transcription factor occupancy and methylation patterns at *trans-*DMRs. Finally, extending these analyses to include other great apes as an outgroup would provide phylogenetic resolution to distinguish changes that occurred in the human vs. the chimpanzee lineage.

Collectively, our findings provide a framework for understanding how DNA methylation divergence contributes to human-specific traits. We demonstrate that while both *cis*- and *trans*-regulatory mechanisms shape interspecies methylation differences, *cis*-acting factors predominate, thus directly linking genomic sequence variation and epigenetic divergence. The coordinated nature of many independent methylation-expression changes across phenotypically relevant pathways is consistent with epigenetic modifications being important targets of natural selection in human evolution, potentially contributing to the regulatory innovations underlying human-chimpanzee phenotypic differences.

## Materials and Methods

### Cell culture and differentiation

We utilized two previously characterized human-chimpanzee hybrid iPSC lines (Hy1-30 and Hy1-25) generated through in vitro cell fusion^23^. Human (two individuals, H20961 and H21976), chimpanzee (two individuals, C3649 and C3647) and human-chimpanzee hybrid induced pluripotent stem cells (iPSCs) were maintained on matrigel in mTeSR1 or mTeSR Plus medium (Stem Cell Technologies cat #85850 or cat #100–0276)^23^. These lines were confirmed to be free of mycoplasma contamination and their identity was confirmed using the RNA sequencing data generated in this study. The Columbia Stem Cell Core Facility differentiated the iPSCs into multiple cell types including dopaminergic neurons (DA), skeletal myocytes (SKM), and hepatocytes (HEP) using published protocols^26,45,95–103^. Cranial neural crest cells were differentiated as described^24^. Primary dental pulp stem cells (DPSC) samples were obtained from two human and two chimpanzee donors as described^24^.

### RNA-sequencing and allele-specific expression analysis

All samples were cryopreserved in liquid nitrogen before RNA extraction^104^. Cells were gently thawed and then washed with PBS and cell pellets were collected via centrifugation at 1000 RPM for 5 min. RNA extraction was performed using the RNeasy Mini Kit (Qiagen, 74104) and DNase digestion was performed using RNase-Free DNase Set (Qiagen, 79254) following standard protocols. RNA quality was assessed using the Agilent Bioanalyzer RNA Pico assay. Only samples with an RNA integrity number (RIN) greater than or equal to 7 were used to prepare cDNA libraries. Libraries were prepared using TruSeq Stranded mRNA kits (Illumina, 20020594) and sequenced on Illumina HiSeq 4000 to generate paired-end reads. An allele-specific expression pipeline adapted from our previous work was implemented, using dual reference genome alignment to hg38 and pantro6 to eliminate mapping bias^23,24^. Reads were processed with SeqPrep, mapped using STAR, deduplicated with Picard, and corrected for allelic bias using Hornet. DESeq2 was used for differential expression analysis with FDR correction, defining significant allele-specific expression as FDR < 0.05 with consistent results across both reference genomes^23,24,26,105–109^. For details, see “Generation of allele-specific count tables” and “Identifying genes with ASE” in the Methods section in Wang et al. (2024)^26^. For hybrid WGBS samples (DA and CNCC), on average, 48.2%±0.2% of all aligned reads are unassignable using SNPs between human and chimpanzee genome, 25.9%±0.2% of the reads are specific to human, and 25.1%±0.2% of reads are specific to chimpanzee. For hybrid RRBS samples (IPSC, SKM and HEP), on average, 88.4%±0.3% of all aligned reads are unassignable using SNPs between human and chimpanzee genome, 5.3%±0.1% of the reads are specific to human, and 5.3%±0.1% of reads are specific to chimpanzee. The proportion of allele-resolvable reads differs markedly between the two bisulfite assays, and this difference is attributable to fragment length and to the genomic territory each assay interrogates. A read can be assigned to a parental allele only if it spans at least one informative human–chimpanzee SNV. RRBS libraries are generated by MspI digestion and are dominated by short fragments (median aligned length 49–50 bp), and their coverage is concentrated in CpG islands and CpG-dense promoters, which are among the most highly conserved sequences in the genome and therefore carry few interspecies SNVs. Among the 16,845,376 unique CpG positions covered by the unphased hybrid RRBS samples, 65.1% have at least one human–chimpanzee SNV within ±50 bp — the footprint reachable by a typical RRBS read — whereas 90.8% have one within ±150 bp, the footprint of a typical WGBS read pair. Consistent with this, the aligned-length distributions of allele-resolved and unassigned reads are essentially identical in all four WGBS libraries (median 150 bp for both), whereas in all six RRBS libraries allele-resolved reads are substantially longer than unassigned reads (medians 70–75 vs 49–50 bp, respectively).

### Bisulfite-seq library preparation, sequencing and mapping

DNA was extracted and processed, and whole-genome bisulfite sequencing (WGBS) and reduced representation bisulfite sequencing (RRBS) was performed at Admera Health Biopharma Services using Illumina Novaseq 6000. WGBS libraries achieved a mean coverage of 9.68× per CpG site, covering 43.6M unique CpG positions in CNCC (hybrid) and 43.4M in CNCC (parental), 41.9M in DA (hybrid, phased) and 38.8M in DA (parental). RRBS libraries achieved a mean coverage of 5.02× per CpG site, covering 5.0M, 4.8M, and 4.5M unique CpG positions in the phased hybrid HEP, IPSC, and SKM libraries respectively, and 3.3M, 3.4M, and 3.2M in the corresponding parental libraries. After phasing the hybrid samples, the total number of CpG sites (union) of all WGBS libraries is 48,994,320 sites, and the union of all CpG sites of RRBS libraries is 7,289,251. The union of all libraries (including WGBS and RRBS) is 49,049,052. We have also included coverage information as a table in Supplemental File 1. To minimize reference mapping bias in cross-species comparisons, reference genomes were first masked at previously identified human-chimpanzee SNV positions using bedtools maskfasta, reducing the potential for allelic mapping preferences^26,110^. Reference genome preparation was performed using Bismark’s bismark_genome_preparation with Bowtie2 as the aligner, which converts unmethylated cytosines in silico to enable bisulfite-aware alignment^111,112^. All samples were aligned to both masked human (hg38) and chimpanzee (pantro6) reference genomes to assess mapping quality and species-specific biases. Reads were quality-trimmed and adapter sequences removed using Cutadapt and Trim Galore with default parameters for RRBS libraries, including the removal of artificially filled-in cytosines from the 3’ end of reads^113,114^. Trimmed reads were aligned using Bismark, and methylation levels were calculated as the ratio of unconverted cytosines to total coverage at each CpG site^111^. For hybrid samples, SNPsplit was used to assign reads to either the human or chimpanzee allele^115^. Strand information was collapsed for symmetric CpG sites and genomic annotations were lifted between genomic assemblies hg38 and pantro6 using UCSC LiftOver with reciprocal mapping to ensure orthologous regions^116,117^.

### PCA and hierarchical clustering

Fractional methylation of individual CpG sites was estimated using bismark_methylation_extractor from Bismark^111^. CpG sites shared across samples were identified using bedtools intersect^110^. PCA was performed using PCASamples from methylKit across all samples and samples within each cell type^118^. Euclidean distance matrices were computed across all samples for hierarchical clustering analysis, with visualization performed using the pheatmap package^119^. Clustering patterns were assessed to validate cell type identity and characterize species-specific methylation signatures.

### Beta-binomial regulatory classification framework

Methylation data from human (H) and chimpanzee (C) parental and hybrid samples were analyzed using a beta-binomial generalized linear modeling framework to account for overdispersion inherent to bisulfite sequencing count data. The data included measurements for parental H-genome (ParH), parental C-genome (ParC), hybrid H-allele (HybH), and hybrid C-allele (HybC) for each genomic locus. For each genomic locus (site or annotated region), methylated counts were modeled using two complementary beta-binomial GLM models to separately evaluate *cis* and *trans* regulatory effects:

**Model I:** *Cis* effect test with methylation data on hybrid alleles

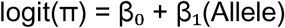

**Model II:** *Trans* effect test using System x Allele interaction term with methylation data in both hybrid and parental alleles

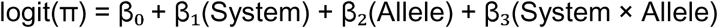

Where π is the probability of methylation, System distinguishes parental from hybrid samples, Allele distinguishes between human and chimpanzee methylation, and the interaction term captures any signals involving the change of allelic differences across generations. Statistical significance was assessed through likelihood ratio tests comparing nested models, testing for system effects, allele effects (*cis*-regulation), and system-by-allele interactions (*trans*-regulation). Genomic loci were classified into five categories: conserved, *trans*-only, *cis*-only, *cis* x *trans* interaction, and *cis* + *trans* effects:

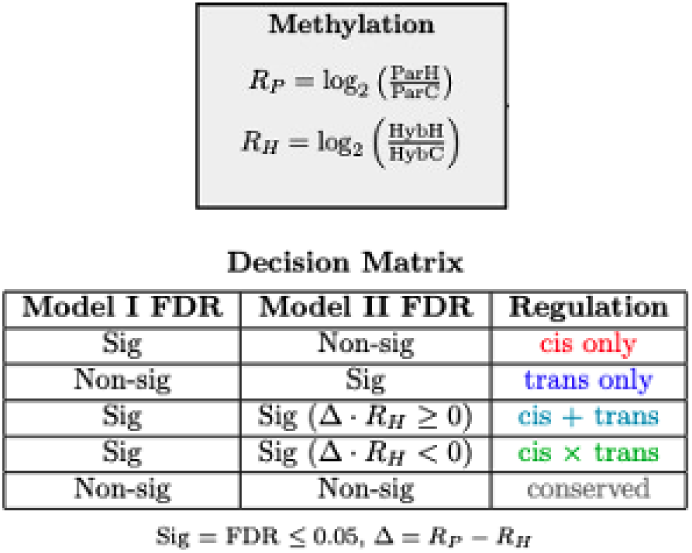

- **Conserved**: No significant allelic differences in either generation (neither parental nor hybrid allelic tests significant)

- ***Trans* only**: Significant allelic difference in parents but not hybrids, indicating trans-acting regulation

- ***Cis* only**: Significant allelic differences in both systems with no significant interaction, indicating consistent cis-acting effects

- ***Cis* × *trans***: Significant allelic differences in both systems with significant interaction, where effects occur in opposite directions

- ***Cis* + *trans***: Significant allelic differences in both systems with significant interaction, where effects are in the same direction

Regional analyses combined CpG-level methylation across defined genomic intervals (for example, promoters). Regional methylation was quantified as pooled fractional methylation, calculated as the total number of methylated cytosines divided by the total coverage across all CpG sites within each region. Nominal p-values < 0.05 from both models were used to provide a schematic overview of site-level CpG methylation in scatterplots and bar plots, preserving the full dynamic range of the data and providing input features for downstream de novo DMR identification. For reporting significant candidate loci under each regulation category as well as formal inference of region-level regulation, Benjamini–Hochberg false discovery rate (FDR) was calculated to account for multiple testing and to evaluate the robustness of inferred relative contribution of regulatory mechanisms across significance thresholds^120^. The same classification and visualization steps were employed using FDR-corrected p-values on loci and regions to confirm that the relative contributions of regulatory classifications were not driven by significance cutoffs alone (Figure 2—figure supplement 1 and 2). To assess the robustness of *cis*-over-*trans* contributions to methylation with respect to the significance cutoffs, the proportion of significant loci and regions assigned to each regulatory category was estimated across a range of FDR cutoffs (FDR < 0.01, 0.05, 0.1, 0.2, 0.25), confirming that the observed *cis* dominance over other divergent regulation categories (including *trans*) was robust to the choice of cutoff (Figure 2—figure supplement 1 and 2).

### Changepoint detection for identifying differentially methylated regions with homogeneous cis- or trans-acting methylation divergence

To identify clusters of CpG sites in the same regulation category, we employed the Pruned Exact Linear Time (PELT) algorithm implemented in the changepoint R package to detect shifts in mean methylation scores within each chromosome segment while accounting for a manually specified penalty against over-fitting^121–123^. We adapted this method to identify regions in the genome where neighboring CpG sites are consistently classified by the previously mentioned beta-binomial model to fall under the same regulatory classification. We first segmented the entire genome into large regions where a break is assigned whenever neighboring CpG sites are more than 5000 base pairs apart. Then, we transform all CpG sites in a segment into binary sequences, where the focal regulation category is 1 and alternative categories are 0. To identify *trans-*DMRs, we set focal pure *trans* regulation to 1, and all other regulation categories to 0. To identify *cis-*DMRs, we set focal pure *cis* regulation to 1, and all other regulation categories to 0. To identify *trans-*DMRs specifically for HOMER enrichment inputs, we used a relaxed gradient scoring approach. We set focal pure *trans* regulation to 1, conserved to 0.75, additive (*cis* + *trans*) and interactive (*cis* x *trans*) regulation to 0.5 and pure *cis* regulation category to 0. To determine the optimal penalty parameter, we applied the CROPS (Changepoints for a Range Of Penalties) method, testing penalty values ranging from 0.01 to 2.0. For each penalty range giving the same number of segments, we recorded the number of detected changepoints and corresponding model fit statistics. The optimal penalty for each segment was determined using elbow point detection via the R package kneedle, which identifies the point of maximum distance from line connecting endpoints in the penalty-versus-changepoints curve^124^. We then applied a homogeneity filter requiring segments to have a minimum target proportion of 0.6, where the target proportion is the fraction of CpG sites within the segment that (i) belong to the target regulatory class under the beta binomial cis/trans classification and (ii) share a consistent species bias in methylation (all human biased or all chimpanzee biased), to ensure biological relevance of identified regions. For segments where fewer than three distinct penalty values yielded different numbers of changepoints, we selected the penalty that maximized the number of segments passing the homogeneity threshold. Segments without detected changepoints were evaluated directly against the homogeneity criterion to assess their potential as single-segment DMRs.

To refine segment boundaries and remove spurious edge effects before merging overlapping DMRs, we implemented a post-processing trimming step that removes segment edges containing sites with regulatory patterns inconsistent with the dominant signal within each segment. This ensures that identified DMRs represent coherent regions of coordinated methylation changes rather than artifacts of the segmentation process. *cis*-DMRs, *trans*-DMRs, *cis* x *trans* and *cis* + *trans* DMRs were identified independently by applying the changepoint detection method separately to each regulatory class.

### Identification of cell-type specific DMRs, species-specific DMRs, and cell-type specific species-specific DMRs

Examples of cell-type specific DMRs (*HOXA9*), species-specific DMRs (*CTSF*), and cell type-specific species-specific DMRs (*LGALS8*) were identified in hybrid samples using the DSS-single (Dispersion Shrinkage for Sequencing data)^125–128^. For cell type-specific DMR identification, we used the formula ∼species + cell type + species:cell type to test for cell type-specific methylation differences, while species-specific DMRs analysis employed ∼species to identify consistent species differences across cell types. Downstream identification of cell-type specific species-specific DMRs identifies the regions where the interaction term species:cell type is significant. The DSS analysis incorporated smoothing across neighboring CpG sites with a smoothing span of 500 bp and required a minimum of 5 CpG sites per region. Statistical significance was determined using Wald tests with FDR < 0.05, and effect size filtering retained regions with methylation difference ≥20%. Integration with gene expression data was performed through Wilcoxon rank-sum tests to identify cell type-specific gene expression patterns associated with cell-type specific DMRs, while permutations (n=1000) were used to assess the significance of coordinated methylation-expression changes in species-specific patterns, which accounts for any background correlation between neighboring genomic features.

### Expression-methylation correlation analysis

Human transcript coordinates and gene body annotations were obtained from the hg38.knownGene.gtf.gz file via UCSC Table Browser^116^, with transcript start positions designated as transcription start sites (TSS). CpG methylation data aggregated within ENCODE-defined promoter regions were analyzed using a beta-binomial classification framework to determine regulatory categories, while corresponding gene expression data underwent parallel hypothesis testing using a negative-binomial model^50,129^. Genes with the same classification for both methylation and expression—either both *cis* or both *trans*—were then analyzed separately. For genes within each regulatory category, the 2 kb surrounding each TSS was split into 10 bins of 200 bp where individual CpG sites were pooled and averaged as fractional methylation values and correlated with expression TPM values. Custom scripts used for binning and correlation are adapted from beta_profile_region.py from CpGtools^130^.

### Sign test selection analysis

To identify coordinated methylation-expression changes potentially under selection, we used a binomial sign test framework that accounts for background probabilities of concordant changes^23,24,26,30,131,132^. Promoter regions were obtained from the ENCODE Project Consortium^129^. Canonical transcripts were downloaded from the UCSC Table Browser hg38 assembly. For genes with multiple annotated promoters, we selected the promoter that is the closest to 5’ UTR of the gene, to ensure each gene is assigned one corresponding promoter^116^. Promoter regions were mapped to canonical transcripts, and functional enrichment using pathways from the Kyoto Encyclopedia of Genes and Genomes (KEGG) and Human Phenotype Ontology (HPO) was assessed using directional sign tests that leverage the independence of *cis*-regulatory regions to detect non-random patterns indicative of lineage-specific selection rather than neutral drift^1,23,24,26,72–76^. For the methylation-only sign test, gene sets with more than 200 genes or fewer than 20 genes with corresponding promoter methylation data were excluded from the analysis. Lineage-specific selection on a gene set was inferred if there was a statistically significant directionality bias in chimpanzee vs. human promoter ASM. For the sign test on genes with putatively repressive methylation—that is, genes where increased methylation within 1 kb of their TSS correspond to decreased expression—we first classified genes as either consistent with repressive methylation (either Ch>Hu methylation paired with Hu>Ch expression, or Hu>Ch methylation paired with Ch>Hu expression) or not consistent. Gene sets with more than 200 genes or fewer than 10 genes with corresponding methylation and expression data were excluded from the analysis. We then reran the sign test using a new background probability defined as the overall fraction of genes with repressive methylation in each cell type. FDRs of the resulting binomial p-values were estimated by randomly permuting the signs of all genes and performing the binomial sign test 10^4^ times. To calculate FDR, we compared each pathway’s empirical binomial p-value against the distribution of permuted p-values. The expected number of false positives was estimated as the average number of pathways reaching that significance level per permutation. FDR was then defined as this expected count of false positives divided by the observed count of pathways with empirical p-values at least as extreme.

### *cis-*DMR sequence variant analysis and *trans-*DMR motif enrichment analysis

High-confidence human-chimpanzee single nucleotide variants (SNVs) were identified using a dual-reference approach with stringent filtering criteria as described previously^26^. Allele-specific methylation within different genomic ranges (25bp, 50bp, 100bp) of CpG-altering variants was calculated. Proportions of regions with Hu>Ch and Hu<Ch methylation were calculated for SNVs in three categories: SNVs that disrupt CpG dinucleotides in humans, SNVs that disrupt CpG sites in chimpanzees, and SNVs that lead to no interspecies changes in CpG content between the species. The significance of the difference in relative proportions of Hu>Ch and Hu<Ch of SNVs in the three categories was assessed using a two-proportion test. For proximity tests, the Mann-Whitney U test^65^ was used to compare the distribution of distances from all *cis* or all *trans-* DMRs to the closest CpG SNV. HOMER’s findMotifsGenome.pl was used to test for motif enrichment in the *trans-*DMR set using the human reference genome hg38 as background ^51^.

### Variance partitioning of methylation divergence

To quantify cis and trans contributions independently of significance thresholds, we partitioned the variance of interspecies methylation divergence, analogous to heritability partitioning. Read counts from CpGs covered in every sample were summed within 1-kb bins, retaining bins with ≥3 CpGs and ≥20 pooled reads per sample. For each bin, we computed three quantities from fractional methylation levels. Total divergence, P = ParH − ParC, is the difference between parental human (ParH) and parental chimpanzee (ParC) cells. The cis component, H = HybH − HybC, is the difference between the human (HybH) and chimpanzee (HybC) alleles within hybrid cells. The trans component, T = P − H, is the part of total divergence not explained by cis. Across bins, the variance (Var) of total divergence decomposes as Var(P) = Var(H) + Var(T) + 2Cov(H,T), where Cov(H,T) is the covariance between cis and trans components; a negative covariance indicates that cis and trans changes tend to act in opposite directions. We report the proportion of variance explained (PVE) by cis, PVE_cis = Var(H)/Var(P), and by trans, PVE_trans = Var(T)/Var(P), together with the covariance term 2Cov(H,T)/Var(P); the three sum to one.

To remove sampling noise, Var(H) was estimated as the covariance between the two hybrid replicates, and Cov(P,H) as the covariance between parental and hybrid estimates, since sampling errors are independent between samples. For CNCC, Var(P) was estimated as the covariance between the two parental replicates. For cell types with one parental line per species, we subtracted the expected sampling variance from Var(P) and report bounds from two corrections. The first uses binomial sampling variances, which underestimate noise. The second scales these by an overdispersion factor (φ), the ratio of the observed to the binomially expected variance between hybrid replicates, which overestimates noise (φ = 1.60–2.71).

Confidence intervals were obtained by genomic block bootstrap (1-Mb blocks, 500 replicates). Chromosomes M and Y were excluded. Sensitivity analyses with ≥50 pooled reads per bin and at single-CpG resolution (≥5 and ≥10 reads) are provided in Supplemental File 1.

## Acknowledgements

We thank Leslie Magtanong for cell culture work and the Columbia Stem Cell Core Facility for performing iPSC differentiation. We are grateful to all members of the Fraser Lab for helpful discussions and feedback on this work. This work was supported by NIH grants R01HG012285 and R35GM156526. Figure panels 1A, 2A and 4A were created with BioRender.

## Data Availability

Raw and processed data generated by this study are publicly available through the Gene Expression Omnibus under accession GSE315270 (BS-seq) and GSE315269 (RNA-seq). The raw and processed bulk RNA-seq data of hybrid and parental iPSCs by Agoglia et al. is available at https://www.ncbi.nlm.nih.gov/geo/query/acc.cgi?acc=GSE144825 and the raw and processed bulk RNA-seq data of hybrid and parental CNCC by Gokhman et al. is available at https://www.ncbi.nlm.nih.gov/geo/query/acc.cgi?acc=GSE146481. The raw and processed bulk RNA-seq data of hybrid hepatocytes and hybrid skeletal myocytes from Wang et al. are available at https://www.ncbi.nlm.nih.gov/geo/query/acc.cgi?acc=GSE232949. scRNA-seq data of parental hepatocytes and parental skeletal myocytes from Barr and Rhodes are available here: https://www.ncbi.nlm.nih.gov/geo/query/acc.cgi?acc=GSE201516. The log fold-changes and associated statistics were used directly from the supplemental material Data S1 of Barr and Rhodes. The human and chimpanzee reference genomes used are available at https://hgdownload.soe.ucsc.edu/goldenPath/hg38/bigZips/ and https://hgdownload.soe.ucsc.edu/goldenPath/panTro6/bigZips/ respectively. The human-chimp pairwise alignment used to identify SNVs and indels is available at https://hgdownload.soe.ucsc.edu/goldenPath/hg38/vsPanTro6/. All scripts for performing analyses and making figures in this manuscript are available at https://github.com/zhenzhenma120/The-causes-and-consequences-of-human-specific-DNA-methylation. All supplemental files are available on Zenodo at https://doi.org/10.5281/zenodo.18205339.

---

**Figure 1—figure supplement 1.**
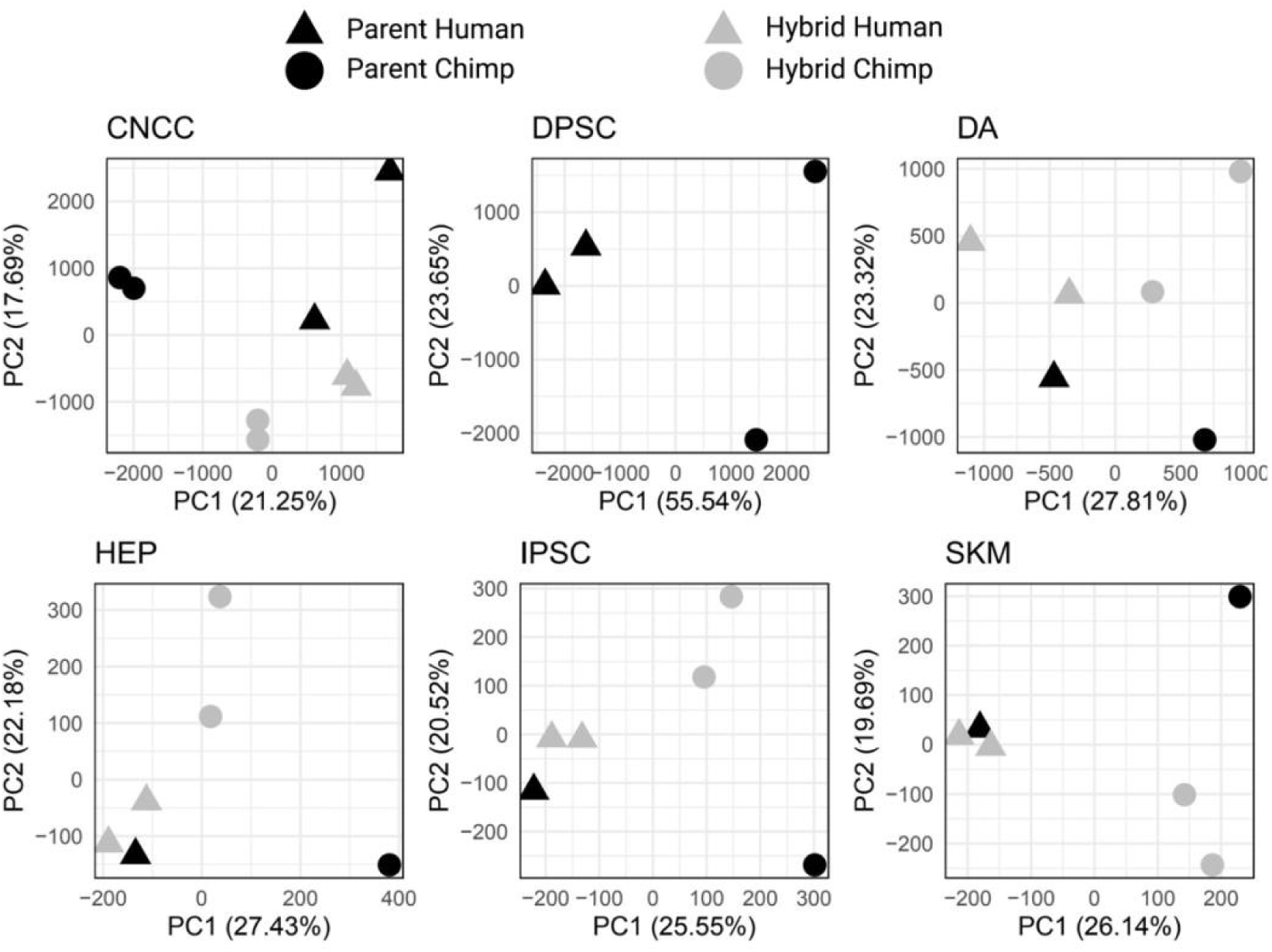
PCA on CpG methylation from BS-seq for individual cell types. Species are clearly separated by PC1, and systems (parent and hybrid) are separated by PC2 in all cell types.

**Figure 1—figure supplement 2.**
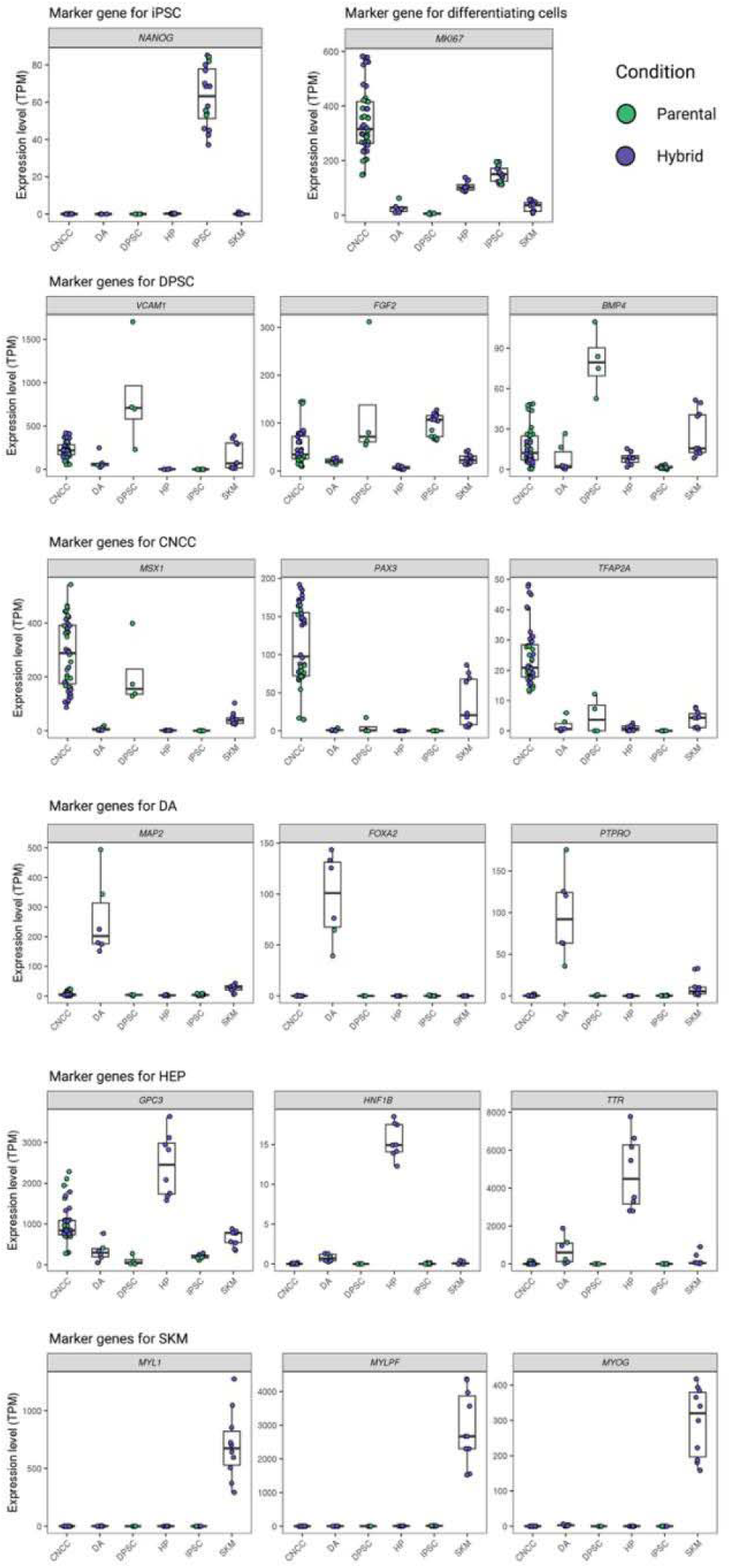
Expression of cell type specific marker genes in different cell types. TPM: transcript per million.

**Figure 2—figure supplement 1.**
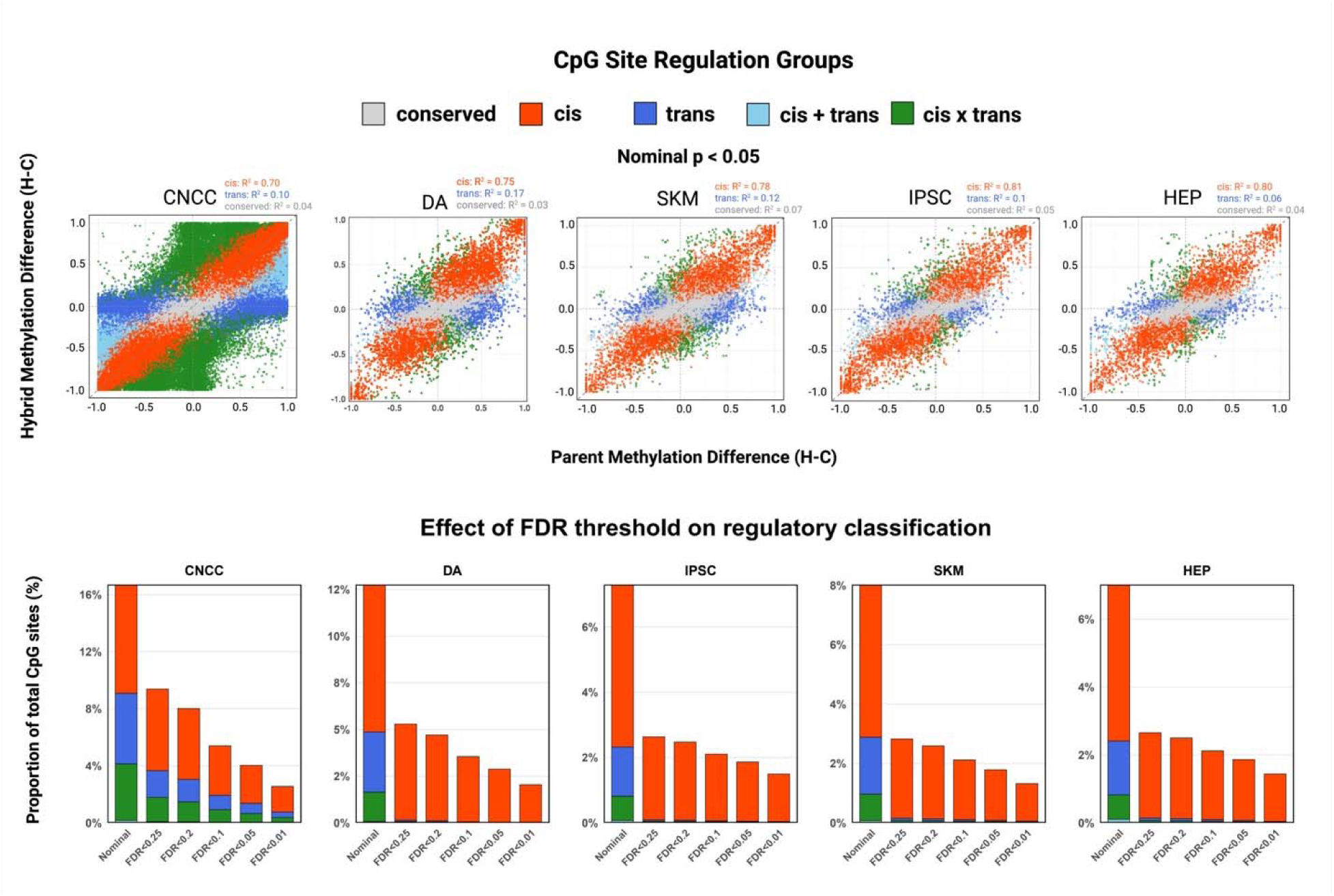
Characterization of regulation groups of all CpG sites for individual cell types. The methylated read counts and unmethylated read counts of each individual CpG site in hybrid and parents is used in the beta-binomial model to classify into regulation groups. Methylation difference is calculated as the difference of fractional methylation between orthologous CpG sites in human and chimpanzee. CNCC has more CpG sites than other cell types due to its high coverage. Scatterplot visualizes CpG site classification at nominal p < 0.05 before FDR correction. After FDR correction, the majority of significant hits are *cis*-regulated CpG sites. Bar plot visualizes the percentages of CpG sites that fall into each regulation group at varying False Discovery Rate (FDR) cutoffs; conserved promoters are used in the calculation but not shown for clarity. For detailed statistics see Supplemental File 1.

**Figure 2—figure supplement 2.**
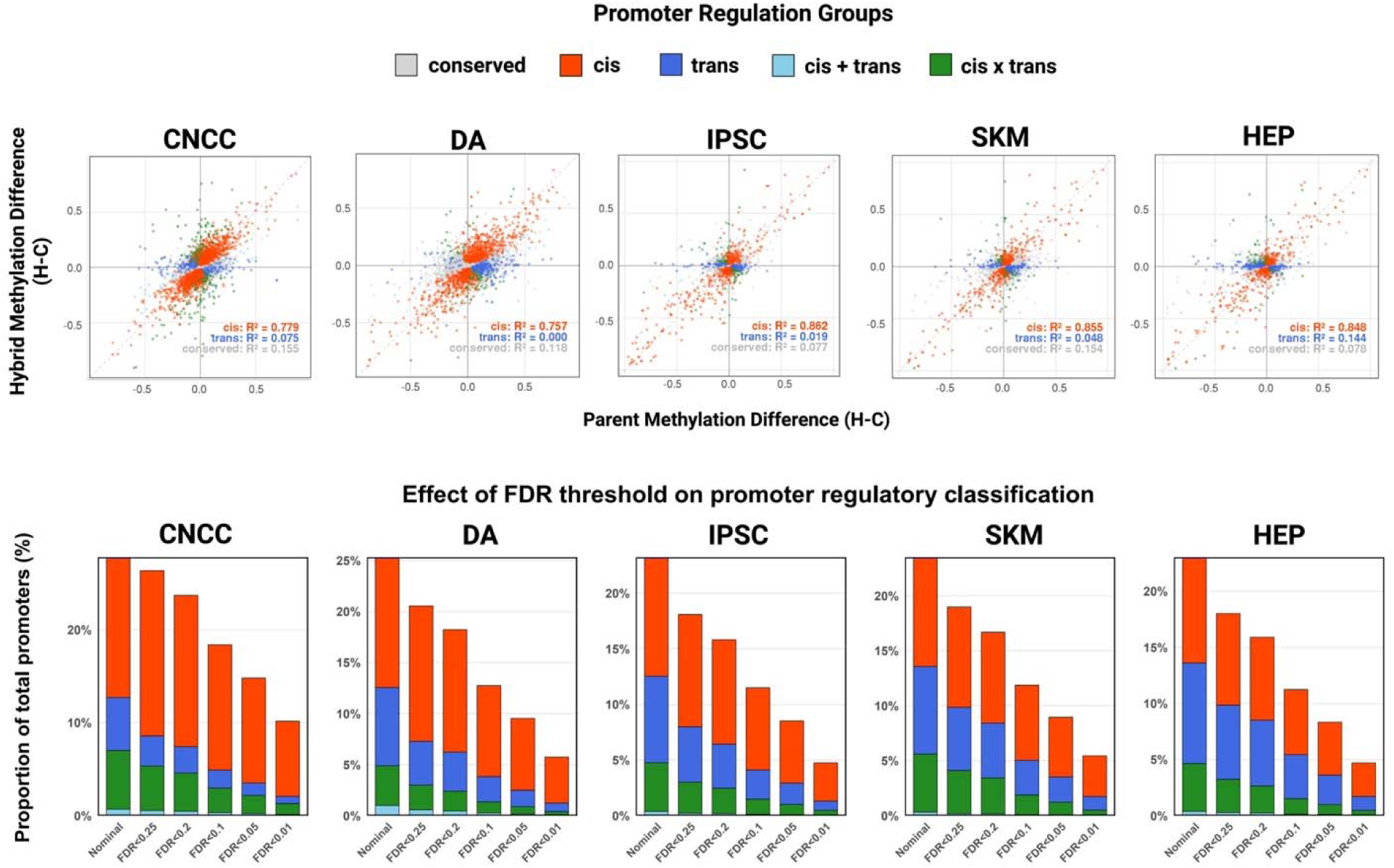
Characterization of regulation groups of all promoters for individual cell types. The pooled methylated and unmethylated read counts in hybrid and parents of each ENCODE promoter region is used in the beta-binomial model to classify into regulation groups. Promoter methylation was quantified as pooled fractional methylation, calculated as the total number of methylated cytosines divided by the total coverage across all CpG sites within each promoter. Scatterplot visualizes promoter classification at FDR < 0.05. Bar plot visualizes the percentages of promoters that fall into each regulation group at varying FDR cutoffs; conserved promoters are used in the calculation but not shown for clarity. See Supplemental File 2 for detailed statistics.

**Figure 2—figure supplement 3.**
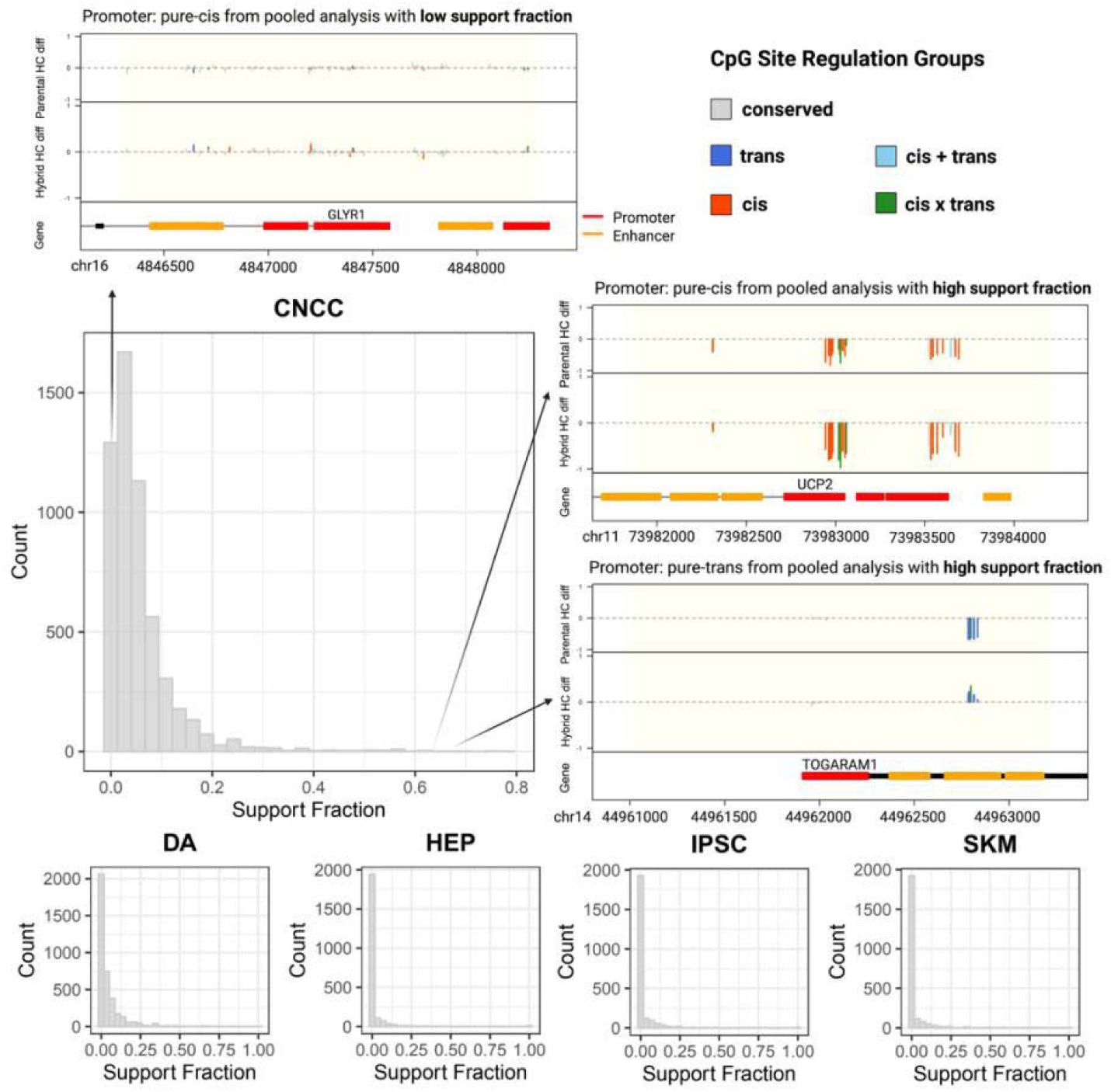
Pooling CpG sites across large genomic regions introduces noise and masks true regulatory patterns. The distribution of support fractions of non-conserved ENCODE-annotated promoters for all cell types showed that most promoter regions have heterogeneous classifications at single CpG site resolution (support fraction: fraction of individual CpG sites out of all sites with the same regulation classification as pooled promoter methylation). Within homogeneous promoters (promoters with high support fractions like *UCP2* and *TOGARAM1*), the majority of CpG sites have the same classification as the promoter. Within heterogeneous promoters (promoters with low support fractions like *GLYR1*), the majority of CpG sites have alternative classifications. Increasing region size results in the aggregation of multiple regulatory units with regions lacking coherent regulatory clusters, which leads to underestimation of differences between contributions of regulatory classifications and masks the true underlying regulatory signal. This highlights the importance of identifying differentially methylated regions (DMRs) exhibiting minimal classification heterogeneity using changepoint detection to prioritize signatures of regulation. See Supplemental File 2 for detailed statistics.

**Figure 3—figure supplement 1.**
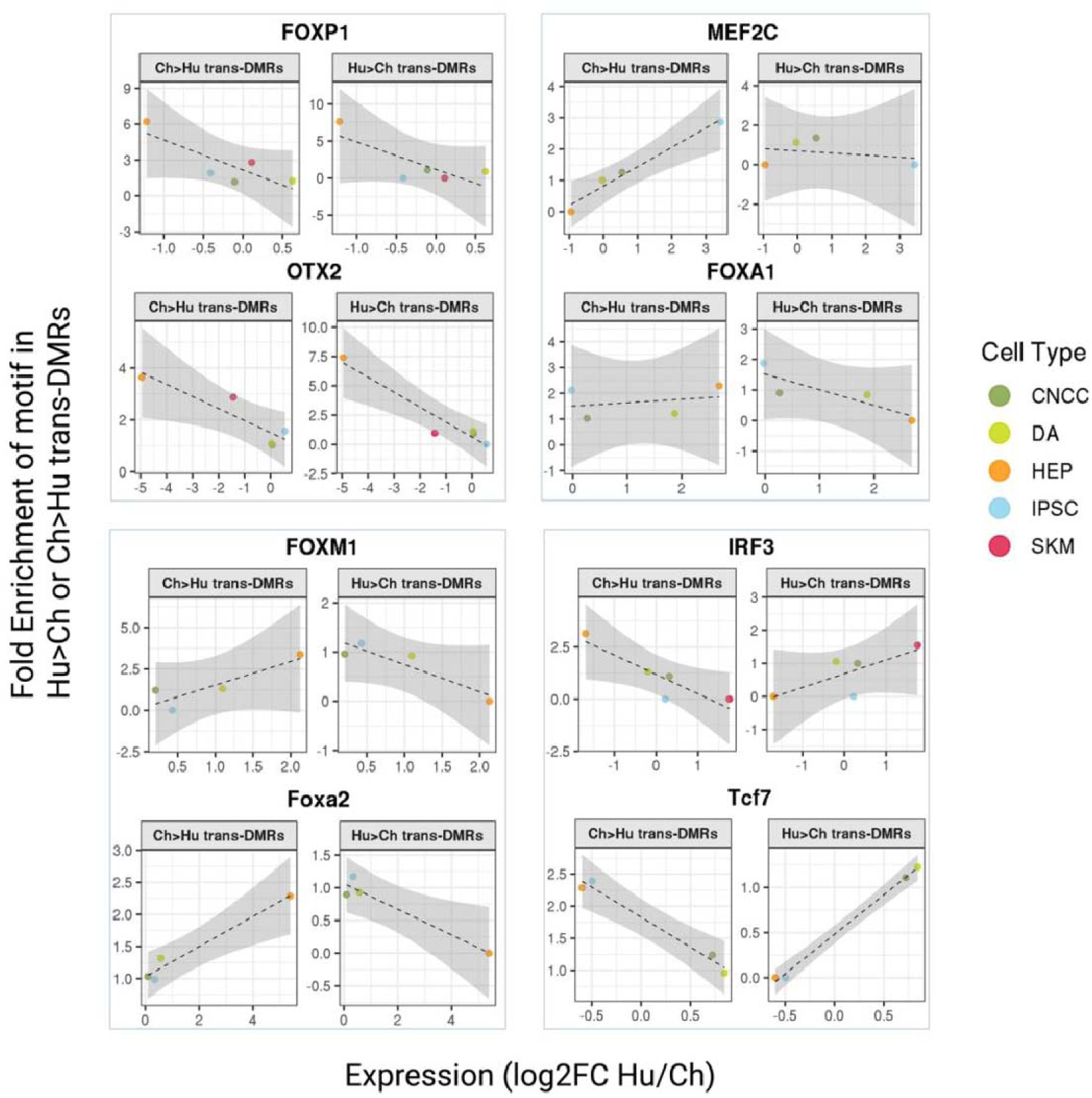
*Trans*-DMR-enriched motifs show different patterns of enrichment relative to differential TF expression. Fold enrichments of motifs in Hu>Ch and Hu<Ch *trans*-DMRs are plotted against differential expression of the matching TF in human vs. chimpanzee parental samples. FOXP1 and OTX2 represent cases where there is no consistent directionality in whether trans-DMRs are human- or chimpanzee-biased when their differential expression increases. MEF2C, FOXA1, FOXM1 and Foxa2 represent cases where higher expression in a species corresponds to enrichment of the motif in regions where methylation is lower in that species, with FOXM1 and Foxa2 showing a stronger pattern. FOXM1 in this figure is the same as Figure 3B. IRF3 and TCF7 represent cases where higher expression in a species corresponds to enrichment of the motif in regions where methylation is higher in that species. See Supplemental File 4 for detailed statistics.

**Figure 3—figure supplement 2.**
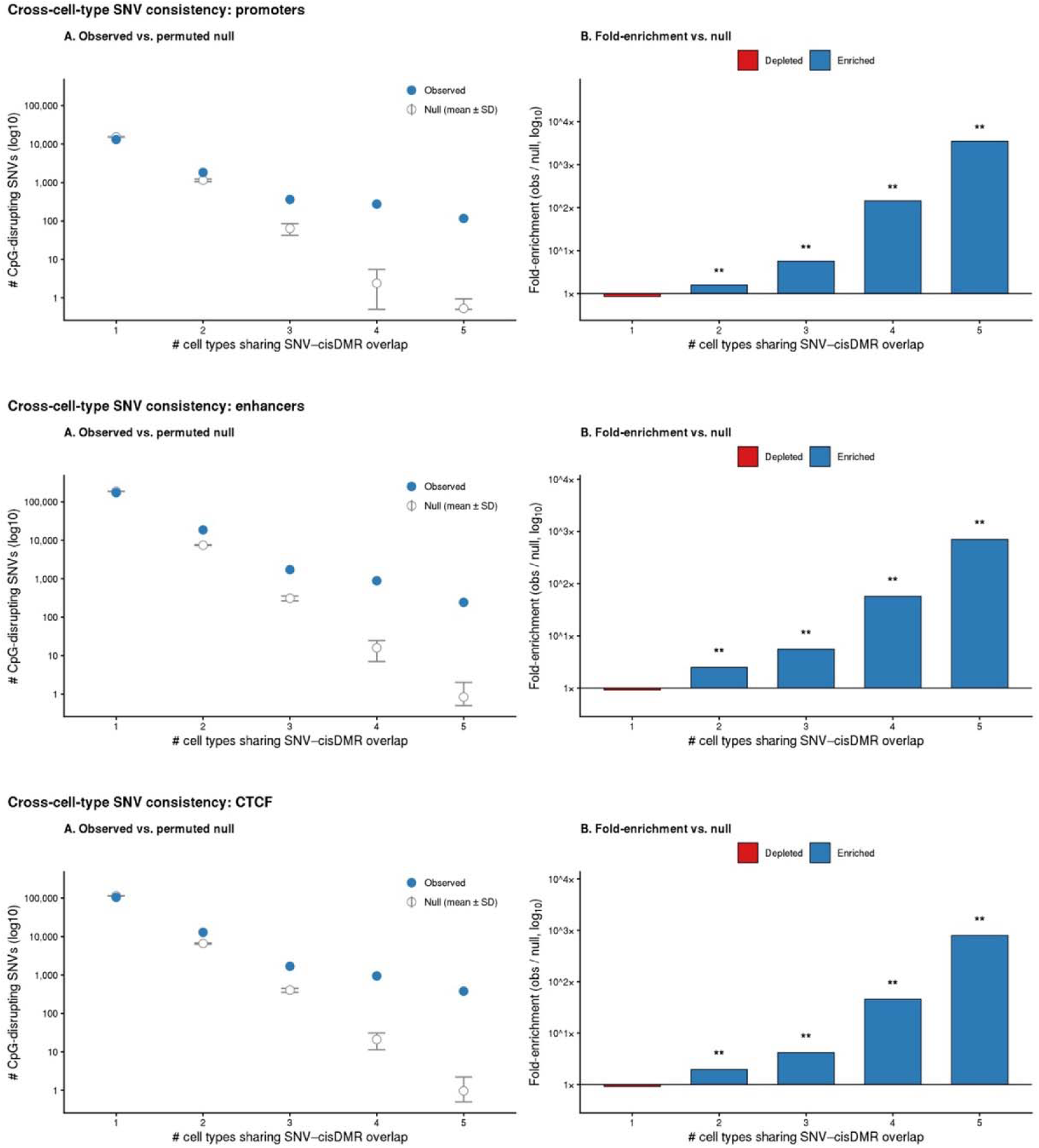
Local methylation effects of CpG SNVs are shared across cell types. For each ENCODE-annotated regulatory region class (promoters, enhancers, CTCF binding sites), CpG SNVs (±100 bp window) were intersected with *cis*-classified DMRs (“pure *cis*” or “*cis+trans*”, BH-corrected FDR < 0.05) across five cell types (CNCC, DA, HEP, IPSC, SKM). For each SNV falling within the tested universe (the union of all assayed regions across cell types), we counted the number of cell types in which it overlapped a cis-DMR. (A) Number of CpG SNVs (log10 scale) sharing cis-DMR overlap across 1–5 cell types. Blue filled points: observed counts; open grey points with error bars: mean ± SD across 1,000 permutations of a null model in which, for each cell type, the same number of regions were randomly sampled from that cell type’s pool of tested-but-conserved regions of the same class. (B) Fold-enrichment of observed over permuted-null counts (log10 scale), with raw fold-change values printed above each bar. Blue bars: enrichment (observed > null); red bars: depletion (observed < null). Asterisks indicate two-sided p-values: *p < 0.05, **p < 0.01, ***p < 0.001 (minimum achievable p = 1/1001 ≈ 0.001 with 1,000 permutations). CpG SNVs were shared across cell types substantially more than expected by chance for all three region classes (mean cell types sharing per SNV-in-cis-DMR: promoters 1.244 vs. null 1.077 ± 0.006; enhancers 1.133 vs. 1.042 ± 0.001; CTCF 1.171 vs. 1.061 ± 0.002; all p < 0.001), with fold-enrichment scaling steeply with the degree of sharing (up to ∼3,500× at promoters, ∼800× at CTCF, and ∼700× at enhancers for SNVs shared across all five cell types). Singleton SNVs (cis-DMR overlap in only one cell type) were correspondingly depleted relative to the null.

**Figure 4—figure supplement 1.**
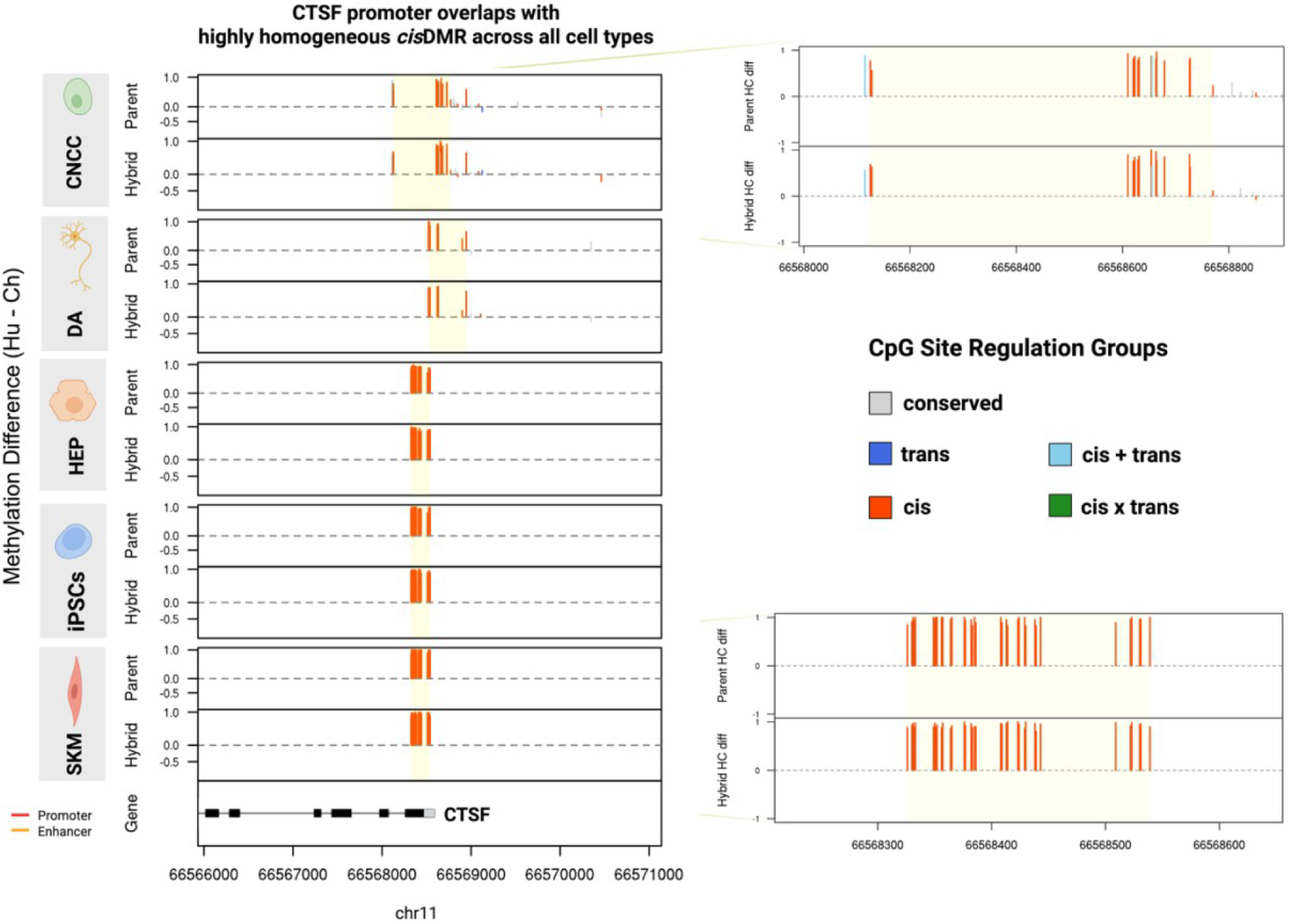
*CTSF* promoter overlaps with highly homogeneous *cis*-DMR across all cell types. Differences of fractional methylation between human and chimpanzee in hybrid and parental cell types are compared. Each vertical line represents methylation difference (Hu-Ch) of a single CpG site, colored by its regulation groups informed by its allele-specific and differential methylation levels. Consistent across all cell types, a homogeneous cluster of cis-regulated CpG sites identified as *cis*-DMR overlaps with the *CTSF* promoter, meaning that besides the entire promoter region being classified as a cis-DMR promoter using pooled methylation counts, the majority of individual CpG sites within the promoter region are individually classified as cis CpG sites. See Supplemental File 3 for detailed statistics.

**Figure 4—figure supplement 2.**
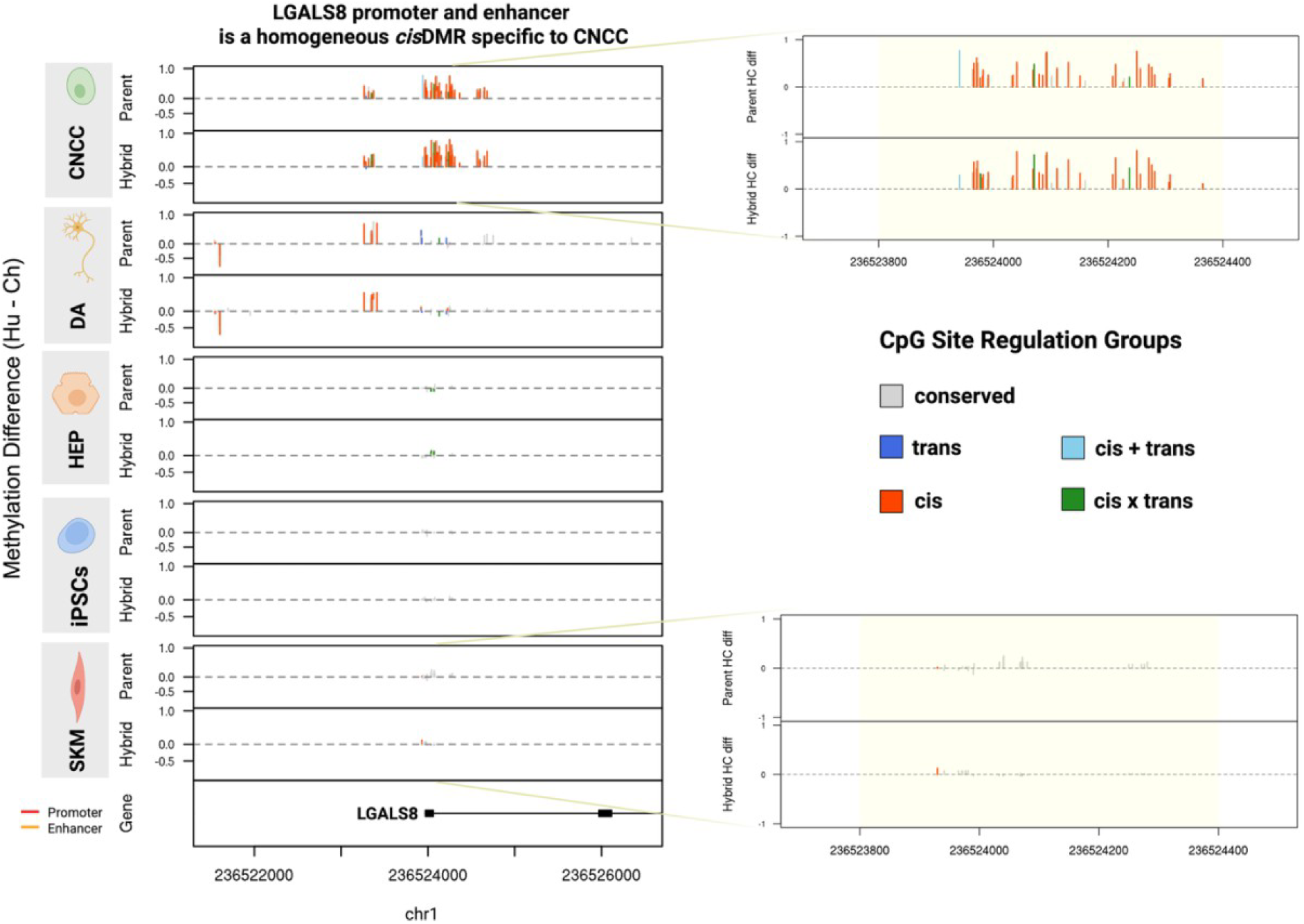
*LGALS8* promoter and enhancer is a homogeneous *cis*-DMR specific to CNCC. Differences of fractional methylation of human and chimpanzee in hybrid and parental cell types are compared. Each vertical line represents methylation difference of a single CpG site, colored by its regulation group. From Figure 4E, it is evident that the *LGALS8* promoter and enhancer region shows allele-specific methylation exclusively in CNCC while showing no allelic differences in other cell types. As seen from this figure, the allele-specific methylated region is also differentially methylated in CNCC parental cells and the majority of individual CpG sites are characterized as cis, meaning that this region is a *cis*-DMR, whereas other cell types show conserved methylation in the same region. See Supplemental File 3 for detailed statistics.

**Figure 4—figure supplement 3.**
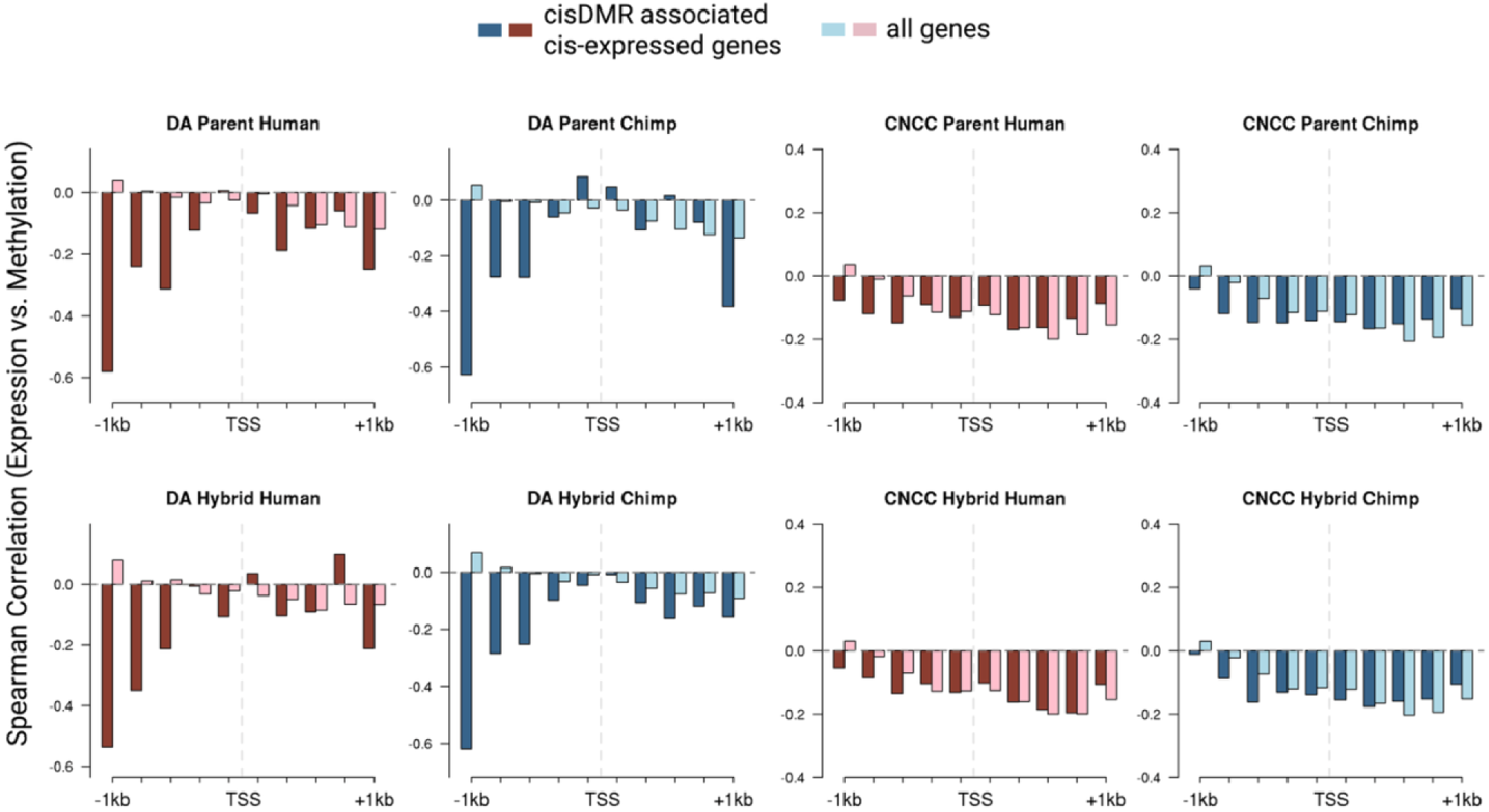
Spearman correlation of expression and methylation for CNCC and DA. Binned expression-methylation correlation profile of genes with both methylation and expression under cis-regulation (dark red and steel blue) and all genes regardless of regulation type (light pink and light blue). Two kb surrounding each transcription start site (TSS) is shown, split into 10 bins of 200 bp where individual CpG sites are pooled and averaged as fractional methylation values and correlated with expression values of the neighboring genes (in TPM). A weak but significant negative correlation is seen between expression and methylation levels near the promoter regions slightly upstream of TSS, consistent with past studies. When restricting to only the genes with cis-regulated methylation and cis-regulated expression, the correlation is stronger, especially in the bin 800 bp - 1 kb upstream of the TSS in DA cells. See Supplemental Files 5 and 6 for significance and detailed statistics.

**Figure 4—figure supplement 4.**
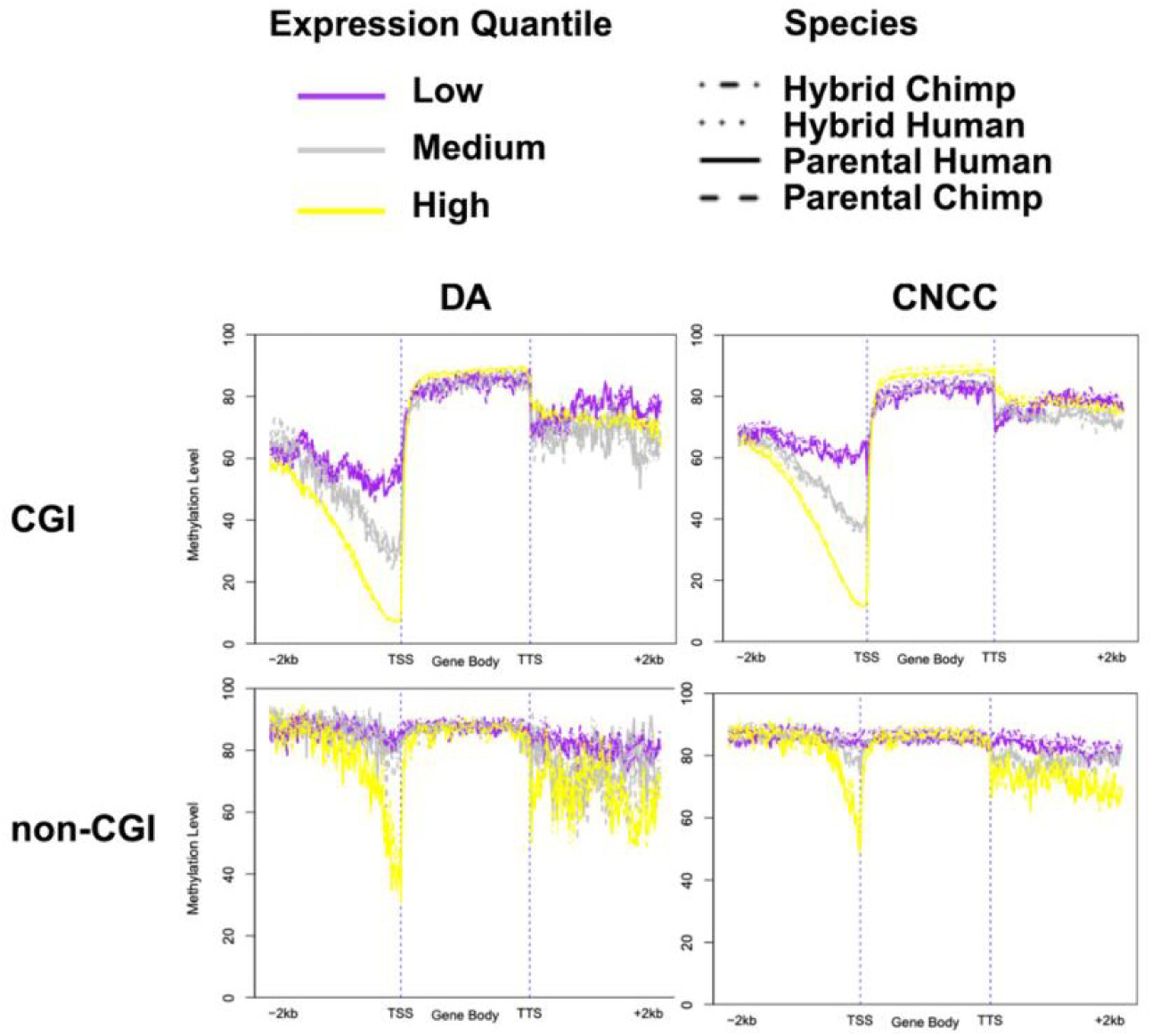
CpG island (CGI) promoters show stronger repressive methylation-expression coupling than non-CGI promoters. Genes were stratified into three equally sized groups based on their expression levels (TPM), and average methylation level of each group is shown in 100 bins across 2000bp upstream of transcription start site (TSS), gene body (all gene bodies scaled to be equal length in this plot), as well as 2000bp downstream of transcription termination site (TTS). A general pattern follows for CGI promoters that high expression is associated with low methylation, medium expression is associated with intermediate methylation levels, and low expression is associated with high methylation, whereas non-CGI promoters shows less stratification. Analysis of both CGI and non-CGI promoters revealed that genes are generally hypomethylated in promoter regions, with highly expressed genes showing a higher degree of hypomethylation in promoters, consistent with the established role of promoter methylation in transcriptional repression. However, CGI promoters showed more pronounced stratification, with highly expressed genes displaying significantly lower promoter methylation levels compared to lowly expressed genes.

**Figure 4—figure supplement 5.**
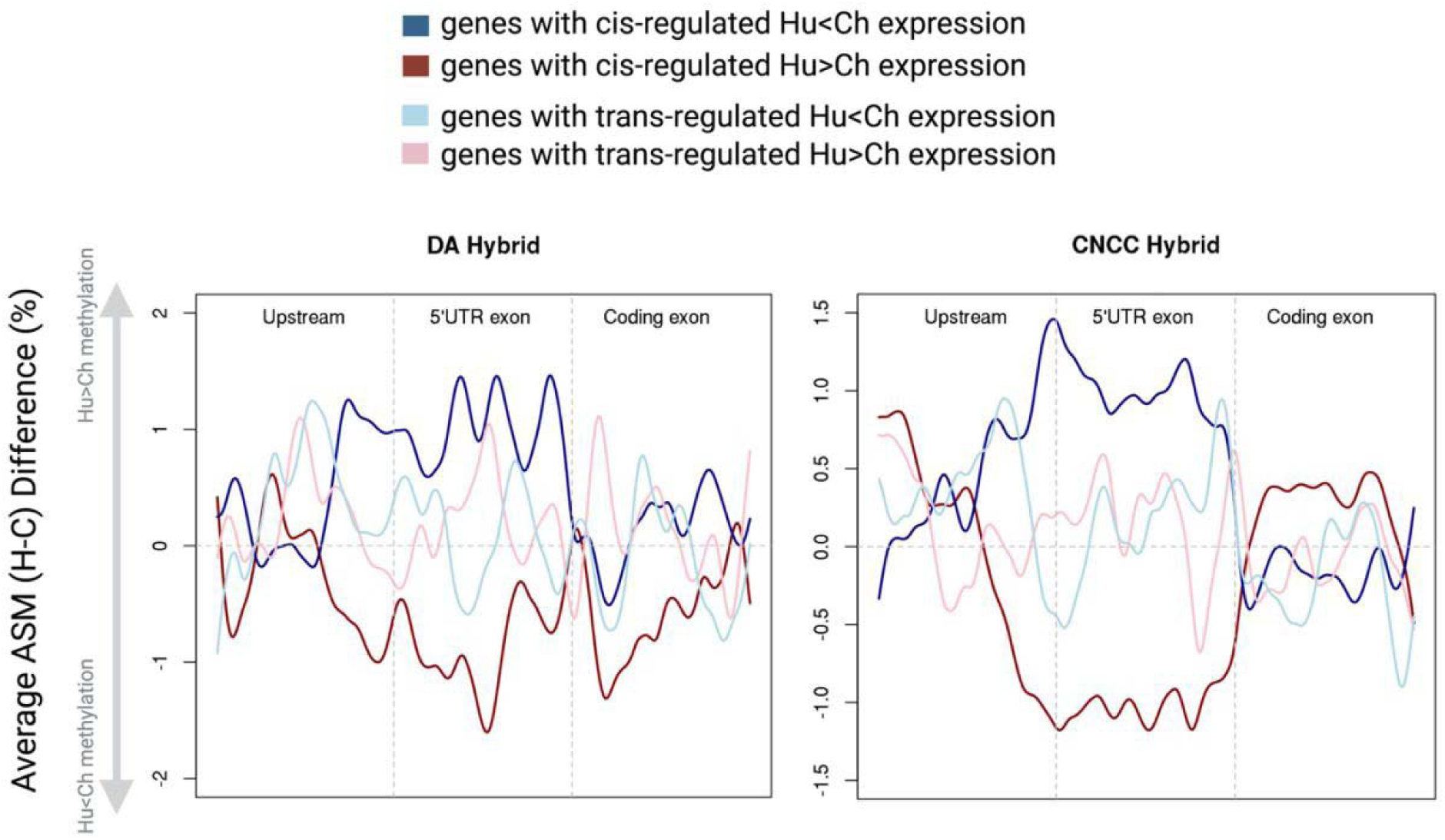
Genes with cis-regulated expression are more likely to exhibit repressive patterns (i.e., higher methylation associated with lower expression) than those with trans-regulated expression. The upstream regions (1000bp upstream of TSS), gene bodies and downstream regions (1000bp downstream of TTS) are each splitted into 100 equal bins where individual CpG sites are pooled and averaged as fractional methylation values. In genes with cis-regulated gene expression, the ones with Hu<CH expression shows Hu>Ch methylation on average across ∼1000bp upstream of and including 5’ UTR region whereas genes with Hu>Ch expression shows Hu<Ch methylation. In genes with trans-regulated gene expression, the repressive pattern is not observed. This is consistent with DNA methylation acting as an upstream cis-regulatory mechanism.

**Figure 5—figure supplement 1.**
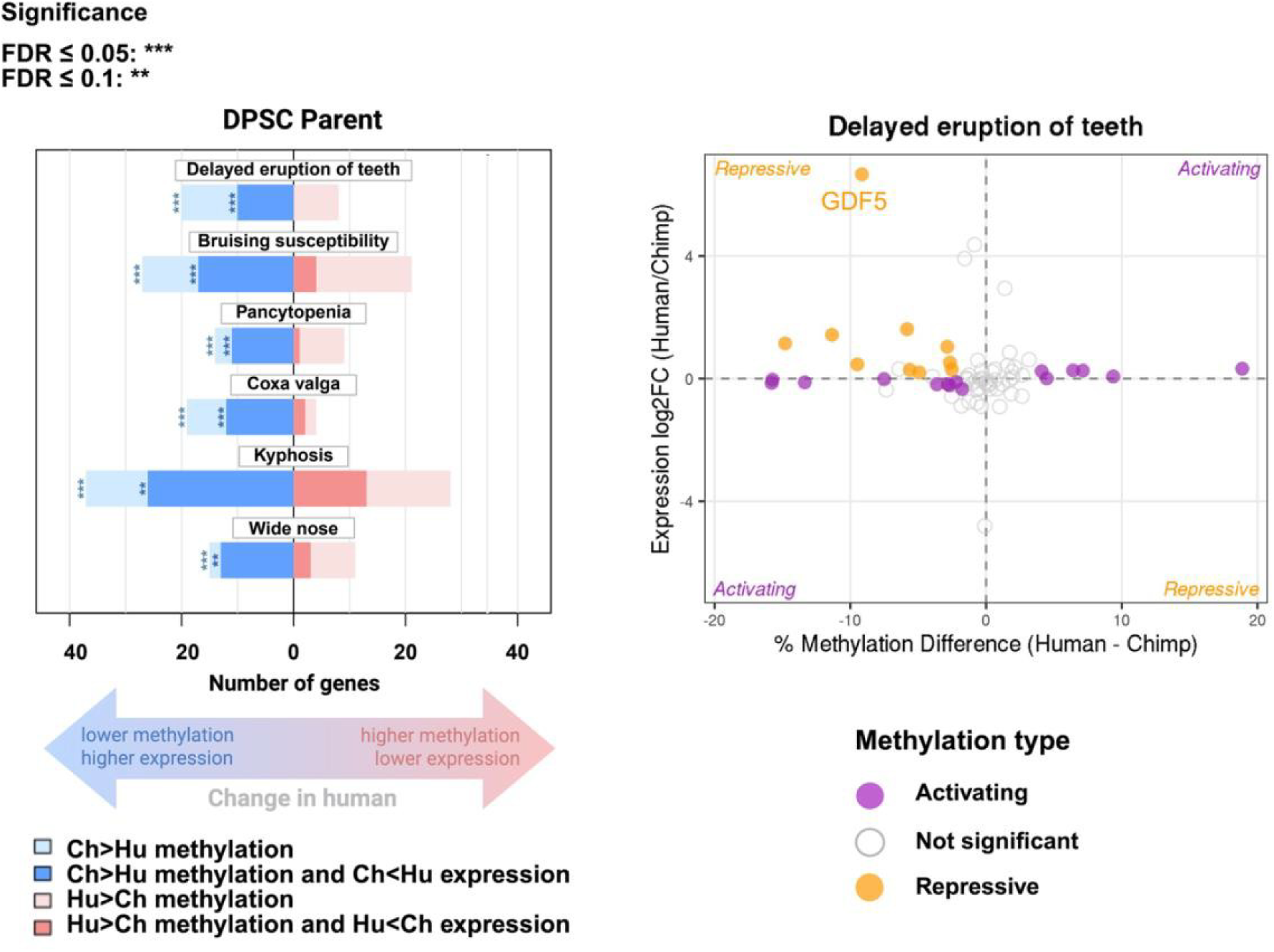
Sign test on DPSC parental methylation and expression data shows significant bias for genes related to delayed eruption of teeth. Left panel: The length of the bars indicates the number of genes in a category with allele-specific expression or methylation in each direction (human or chimpanzee-biased). The bars in bolded colors represent the number of genes within the pathway where higher methylation is associated with repressed gene expression, whereas lighter colors represent genes where higher methylation is associated with increased gene expression. Right panel: Differential expression log fold-changes are plotted against methylation. Notably, in pathway “delayed eruption of teeth”, *GDF5* has >30-fold higher expression and ∼10% lower promoter methylation in human compared to chimpanzee DPSCs. Since the sign test assumes independence of gene expression changes within a pathway, results on parental data (which do not exclude trans-acting factors that may be shared across multiple genes) are suggestive but cannot reject the null hypothesis of neutral evolution. DPSC parental samples had a strong genome-wide human bias in promoter methylation, with 68% of promoters showing human-biased methylation compared to 32% chimp-biased, increasing the significance of gene sets with chimp-biased methylation. See Supplemental File 7 for detailed statistics.

