## Supplemental Figures for "*Cis*-regulation and natural selection in the evolution of human DNA methylation"

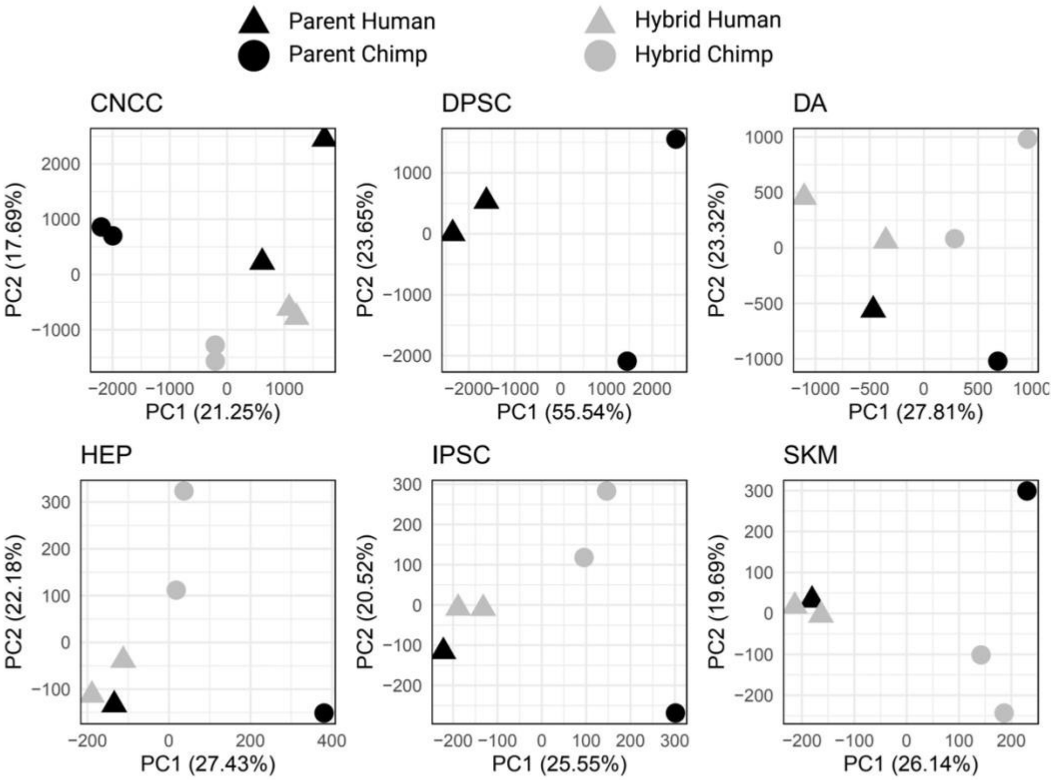


**Figure 1—figure supplement 1. PCA on CpG methylation from BS-seq for individual cell types**. Species are clearly separated by PC1, and systems (parent and hybrid) are separated by PC2 in all cell types.

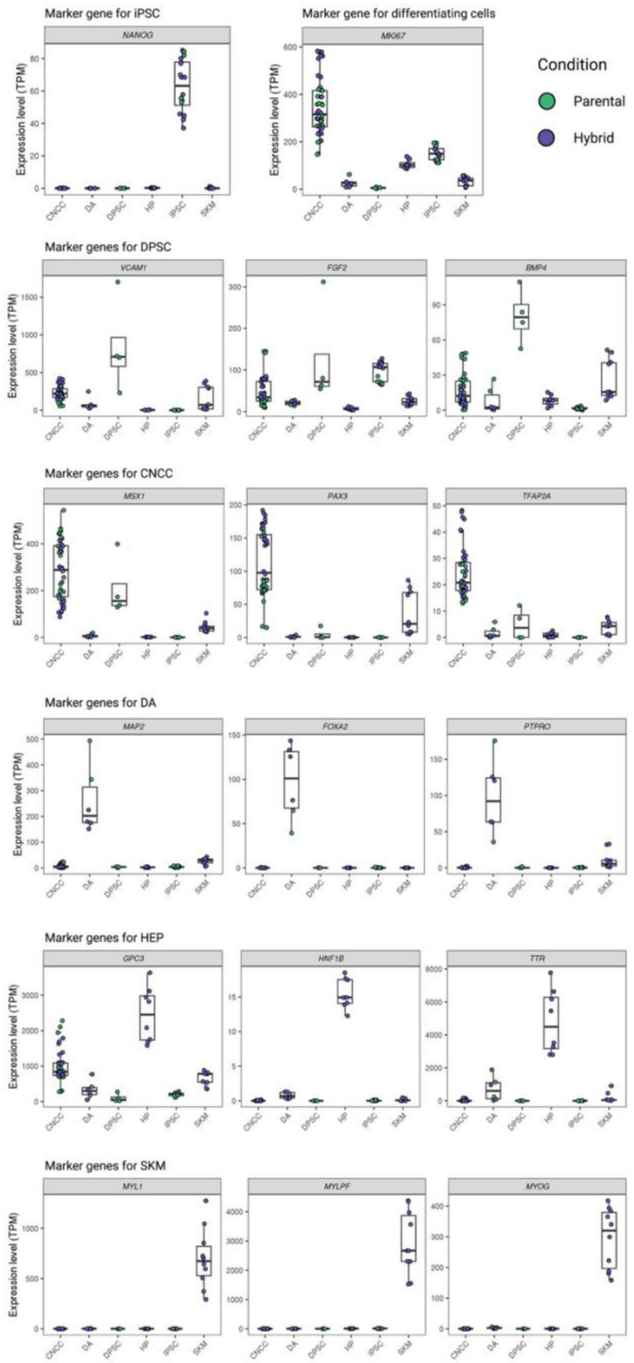


**Figure 1—figure supplement 2. Expression of cell type specific marker genes in different cell types.** TPM: transcript per million.

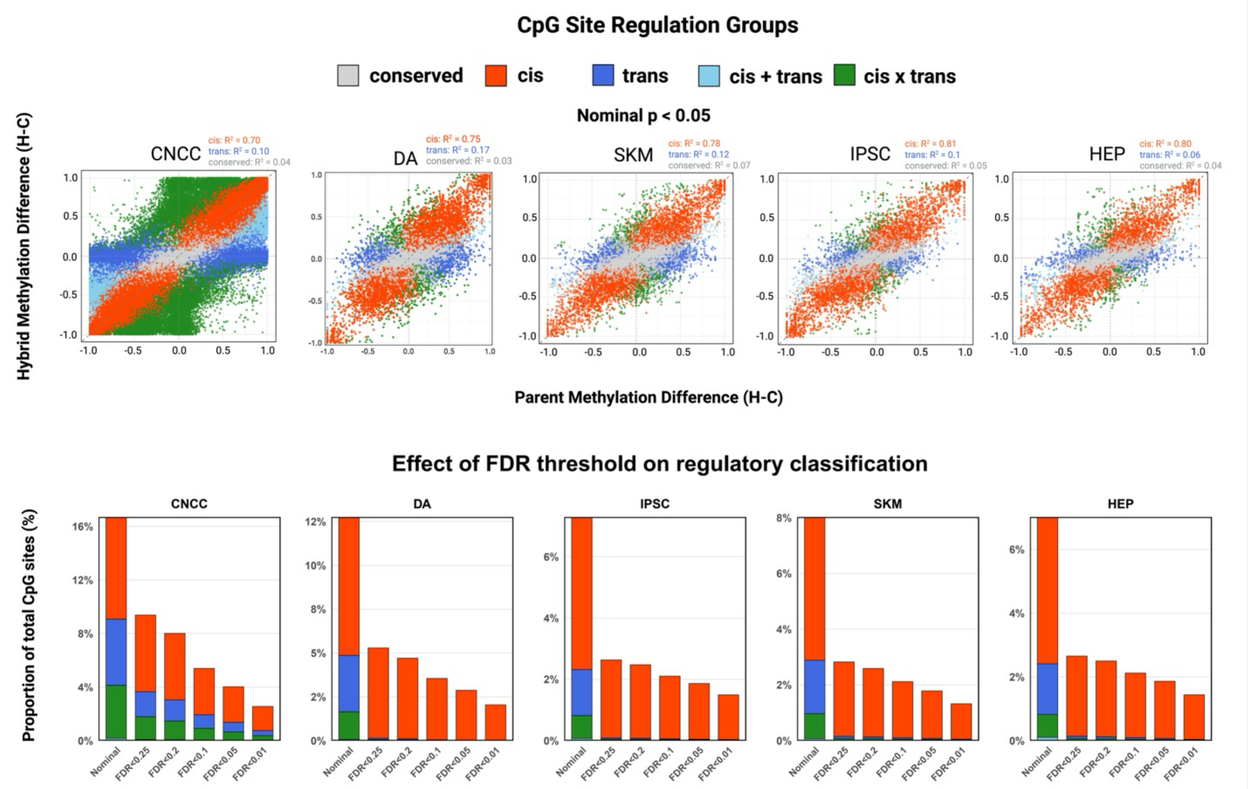


**Figure 2—figure supplement 1. Characterization of regulation groups of all CpG sites for individual cell types.** The methylated read counts and unmethylated read counts of each individual CpG site in hybrid and parents is used in the beta-binomial model to classify into regulation groups. Methylation difference is calculated as the difference of fractional methylation between orthologous CpG sites in human and chimpanzee. CNCC has more CpG sites than other cell types due to its high coverage. Scatterplot visualizes CpG site classification at nominal $p < 0.05$ before FDR correction. After FDR correction, the majority of significant hits are cis-regulated CpG sites. Bar plot visualizes the percentages of CpG sites that fall into each regulation group at varying False Discovery Rate (FDR) cutoffs; conserved promoters are used in the calculation but not shown for clarity. For detailed statistics see Supplemental File 1.

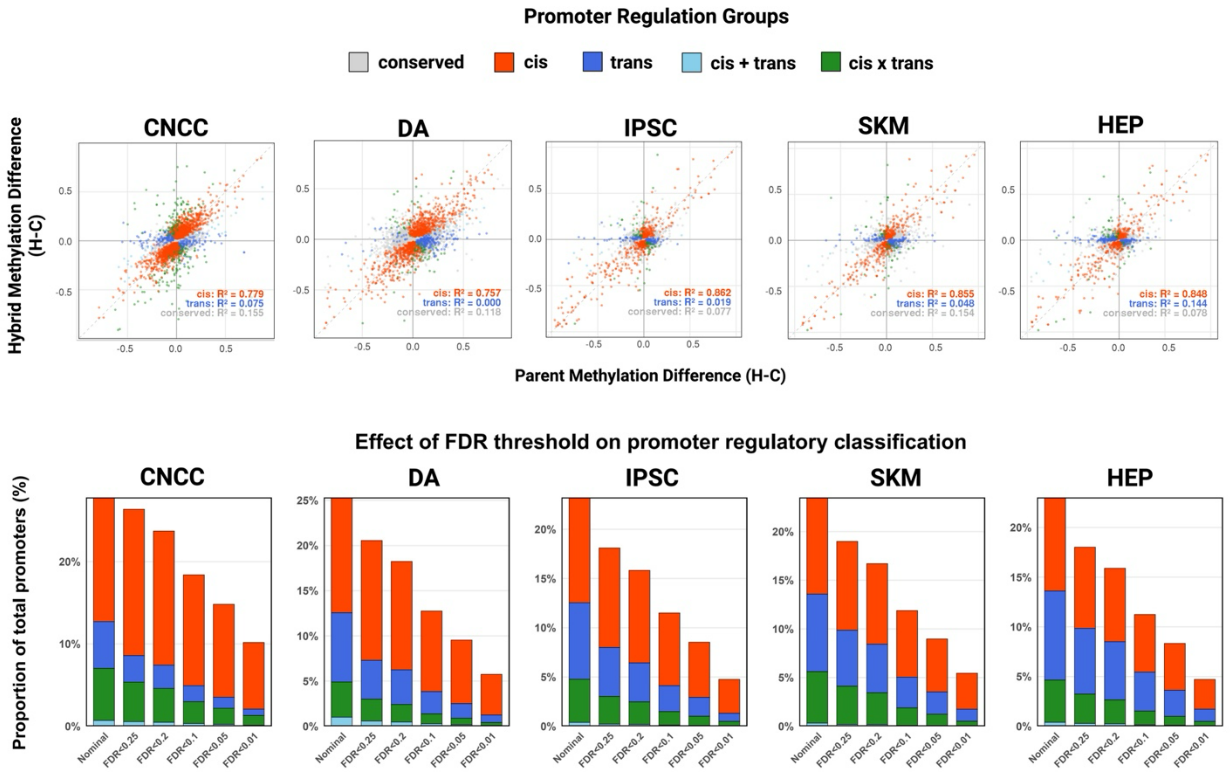


**Figure 2—figure supplement 2. Characterization of regulation groups of all promoters for individual cell types.** The pooled methylated and unmethylated read counts in hybrid and parents of each ENCODE promoter region is used in the beta-binomial model to classify into regulation groups. Promoter methylation was quantified as pooled fractional methylation, calculated as the total number of methylated cytosines divided by the total coverage across all CpG sites within each promoter. Scatterplot visualizes promoter classification at FDR < 0.05. Bar plot visualizes the percentages of promoters that fall into each regulation group at varying FDR cutoffs; conserved promoters are used in the calculation but not shown for clarity. See Supplemental File 2 for detailed statistics.

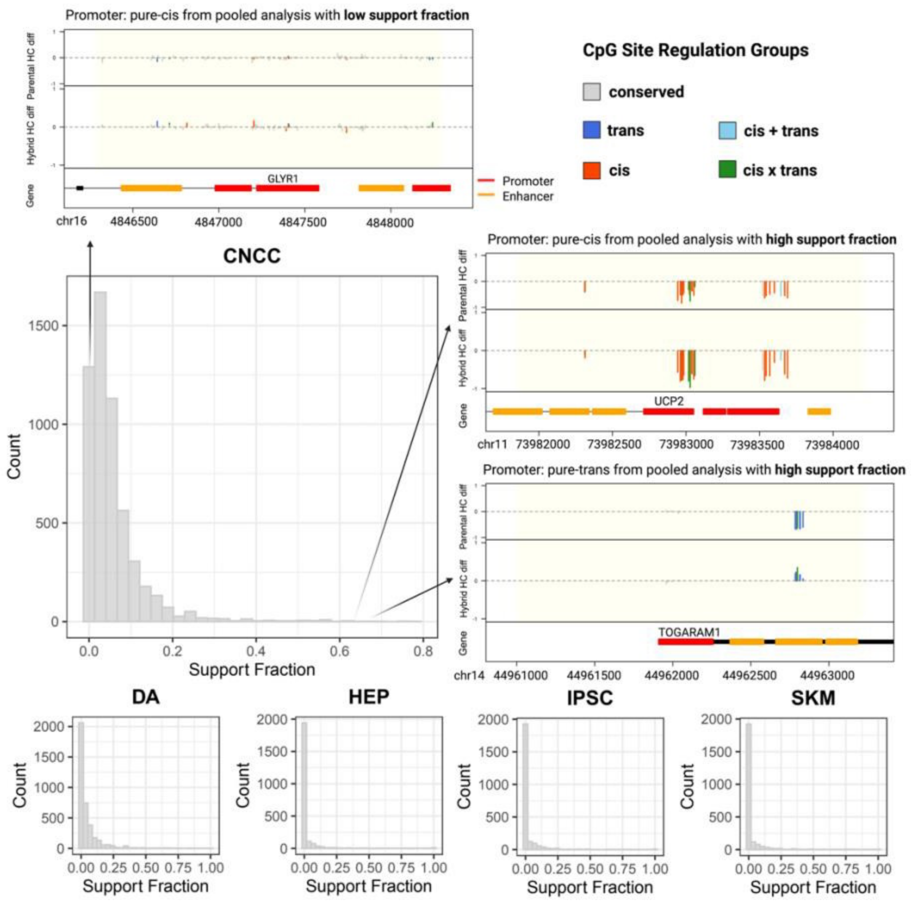


**Figure 2—figure supplement 3. Pooling CpG sites across large genomic regions introduces noise and masks true regulatory patterns.** The distribution of support fractions of non-conserved ENCODE-annotated promoters for all cell types showed that most promoter regions have heterogeneous classifications at single CpG site resolution (support fraction: fraction of individual CpG sites out of all sites with the same regulation classification as pooled promoter methylation). Within homogeneous promoters (promoters with high support fractions like *UCP2* and *TOGARAM1*), the majority of CpG sites have the same classification as the promoter. Within heterogeneous promoters (promoters with low support fractions like *GLYR1*), the majority of CpG sites have alternative classifications. Increasing region size results in the aggregation of multiple regulatory units with regions lacking coherent regulatory clusters, which leads to underestimation of differences between contributions of regulatory classifications and masks the true underlying regulatory signal. This highlights the importance of identifying differentially methylated regions (DMRs) exhibiting minimal classification heterogeneity using changepoint detection to prioritize signatures of regulation. See Supplemental File 2 for detailed statistics.

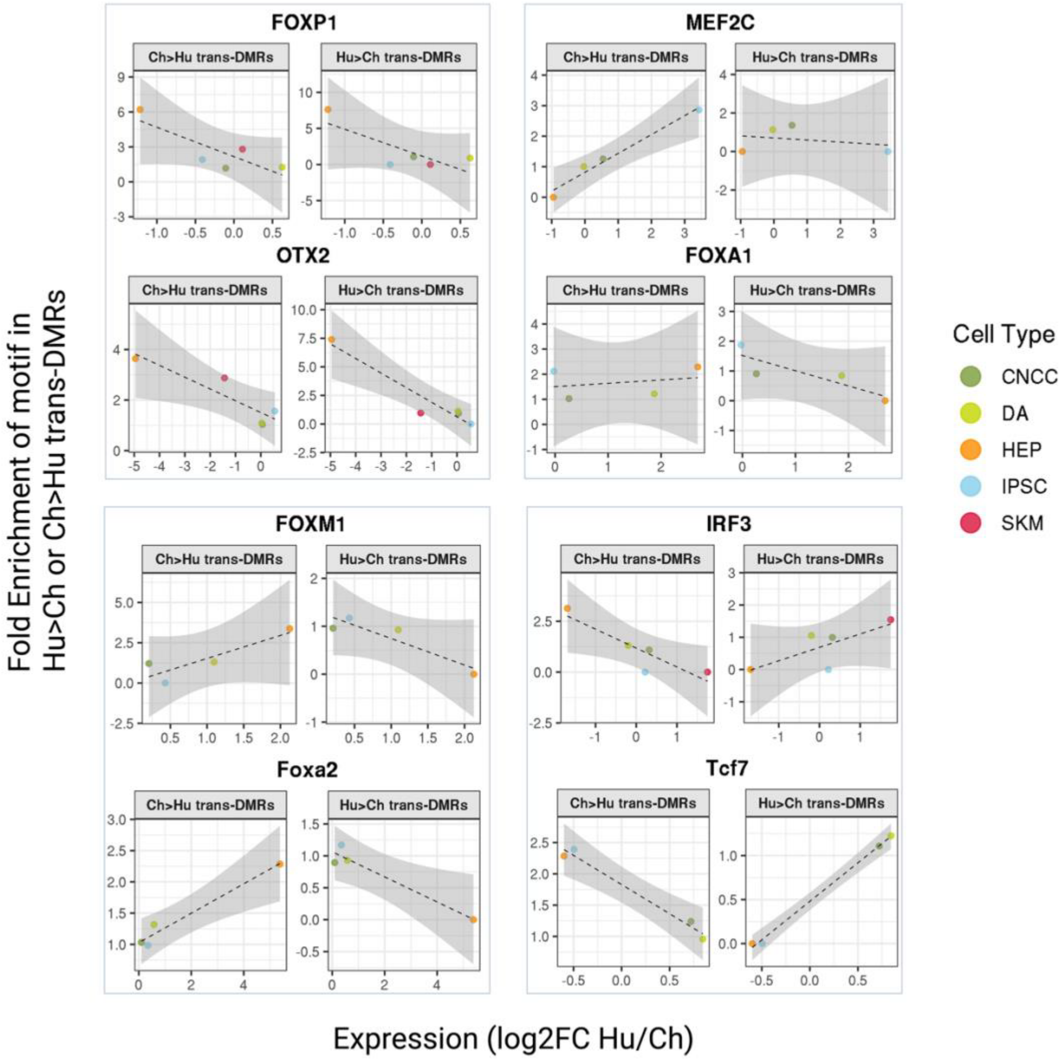


**Figure 3—figure supplement 1. *Trans*-DMR-enriched motifs show different patterns of enrichment relative to differential TF expression.** Fold enrichments of motifs in Hu>Ch and Hu<Ch *trans*-DMRs are plotted against differential expression of the matching TF in human vs. chimpanzee parental samples. FOXP1 and OTX2 represent cases where there is no consistent directionality in whether trans-DMRs are human- or chimpanzee-biased when their differential expression increases. MEF2C, FOXA1, FOXM1 and Foxa2 represent cases where higher expression in a species corresponds to enrichment of the motif in regions where methylation is lower in that species, with FOXM1 and Foxa2 showing a stronger pattern. FOXM1 in this figure is the same as Figure 3B. IRF3 and TCF7 represent cases where higher expression in a species corresponds to enrichment of the motif in regions where methylation is higher in that species. See Supplemental File 4 for detailed statistics.

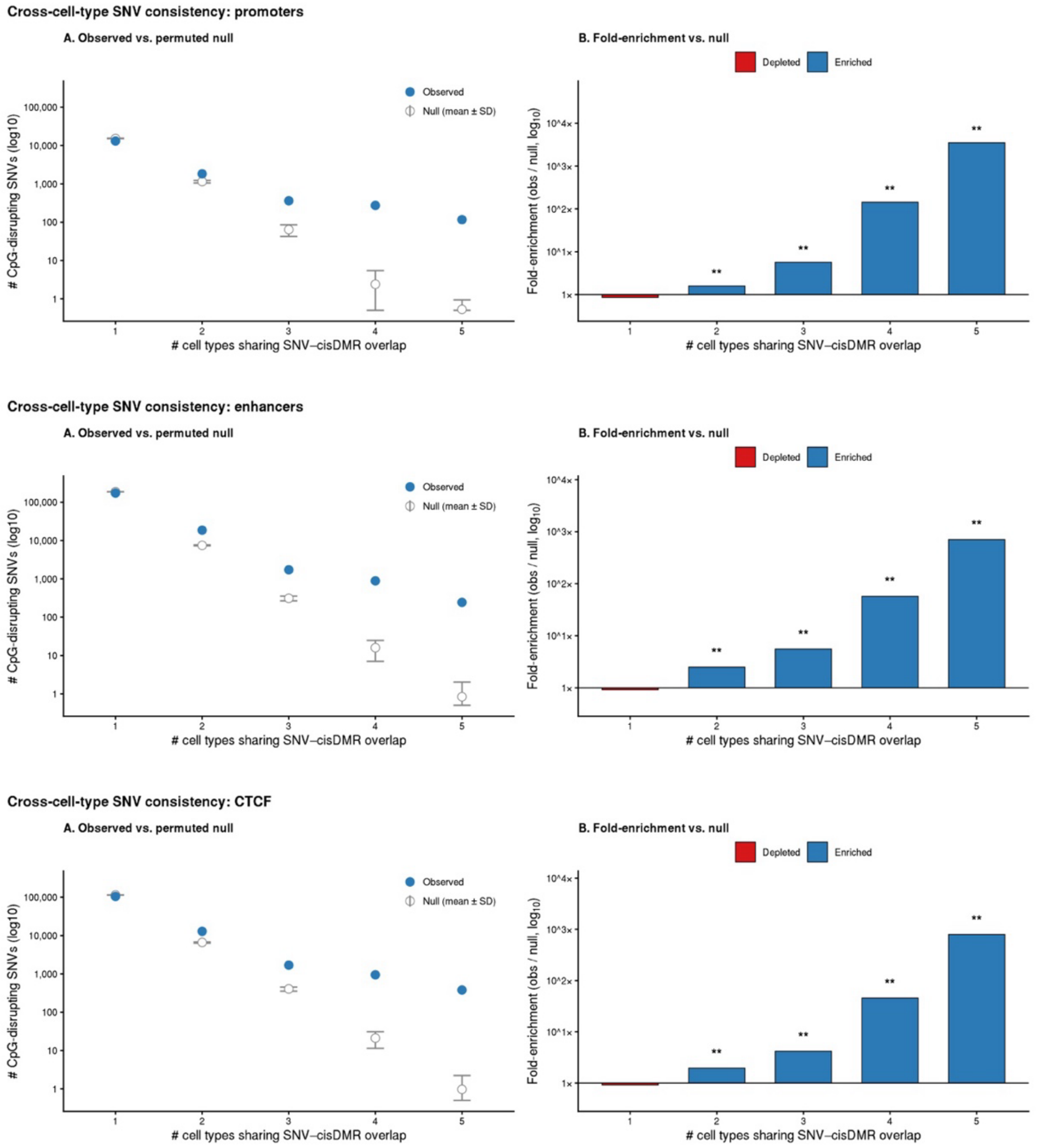


**Figure 3—figure supplement 2. Local methylation effects of CpG SNVs are shared across cell types.** For each ENCODE-annotated regulatory region class (promoters, enhancers, CTCF binding sites), CpG SNVs (±100 bp window) were intersected with cis-classified DMRs ("pure *cis*" or "*cis+trans*", BH-corrected FDR < 0.05) across five cell types (CNCC, DA, HEP, IPSC, SKM). For each SNV falling within the tested universe (the union of all assayed regions across cell types), we counted the number of cell types in which it overlapped a cis-DMR. (A) Number of CpG SNVs (log10 scale) sharing cis-DMR overlap across 1–5 cell types. Blue filled points: observed counts; open grey points with error bars: mean ± SD across 1,000 permutations of a null model in which, for each cell type, the same number of regions were randomly sampled from that cell type's pool of tested-but-conserved regions of the same class. (B) Fold-enrichment of observed over permuted-null counts (log10 scale), with raw

fold-change values printed above each bar. Blue bars: enrichment (observed > null); red bars: depletion (observed < null). Asterisks indicate two-sided p-values: *p < 0.05, **p < 0.01, ***p < 0.001 (minimum achievable p = 1/1001 ≈ 0.001 with 1,000 permutations). CpG SNVs were shared across cell types substantially more than expected by chance for all three region classes (mean cell types sharing per SNV-in-cis-DMR: promoters 1.244 vs. null 1.077 ± 0.006; enhancers 1.133 vs. 1.042 ± 0.001; CTCF 1.171 vs. 1.061 ± 0.002; all p < 0.001), with fold-enrichment scaling steeply with the degree of sharing (up to ~3,500× at promoters, ~800× at CTCF, and ~700× at enhancers for SNVs shared across all five cell types). Singleton SNVs (cis-DMR overlap in only one cell type) were correspondingly depleted relative to the null.

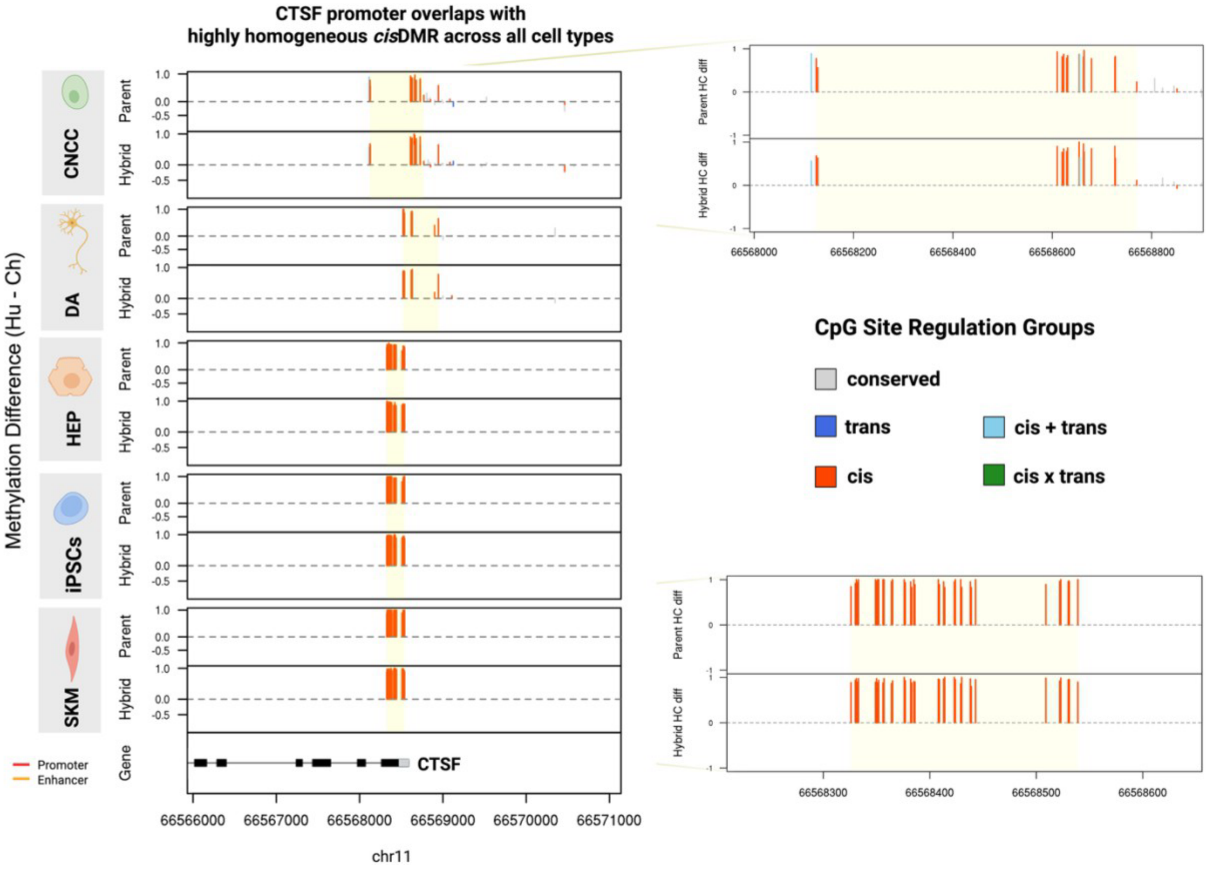


**Figure 4—figure supplement 1.** ***CTSF* promoter overlaps with highly homogeneous *cis*-DMR across all cell types.** Differences of fractional methylation between human and chimpanzee in hybrid and parental cell types are compared. Each vertical line represents methylation difference (Hu-Ch) of a single CpG site, colored by its regulation groups informed by its allele-specific and differential methylation levels. Consistent across all cell types, a homogeneous cluster of cis-regulated CpG sites identified as *cis*-DMR overlaps with the *CTSF* promoter, meaning that besides the entire promoter region being classified as a cis-DMR promoter using pooled methylation counts, the majority of individual CpG sites within the promoter region are individually classified as cis CpG sites. See Supplemental File 3 for detailed statistics.

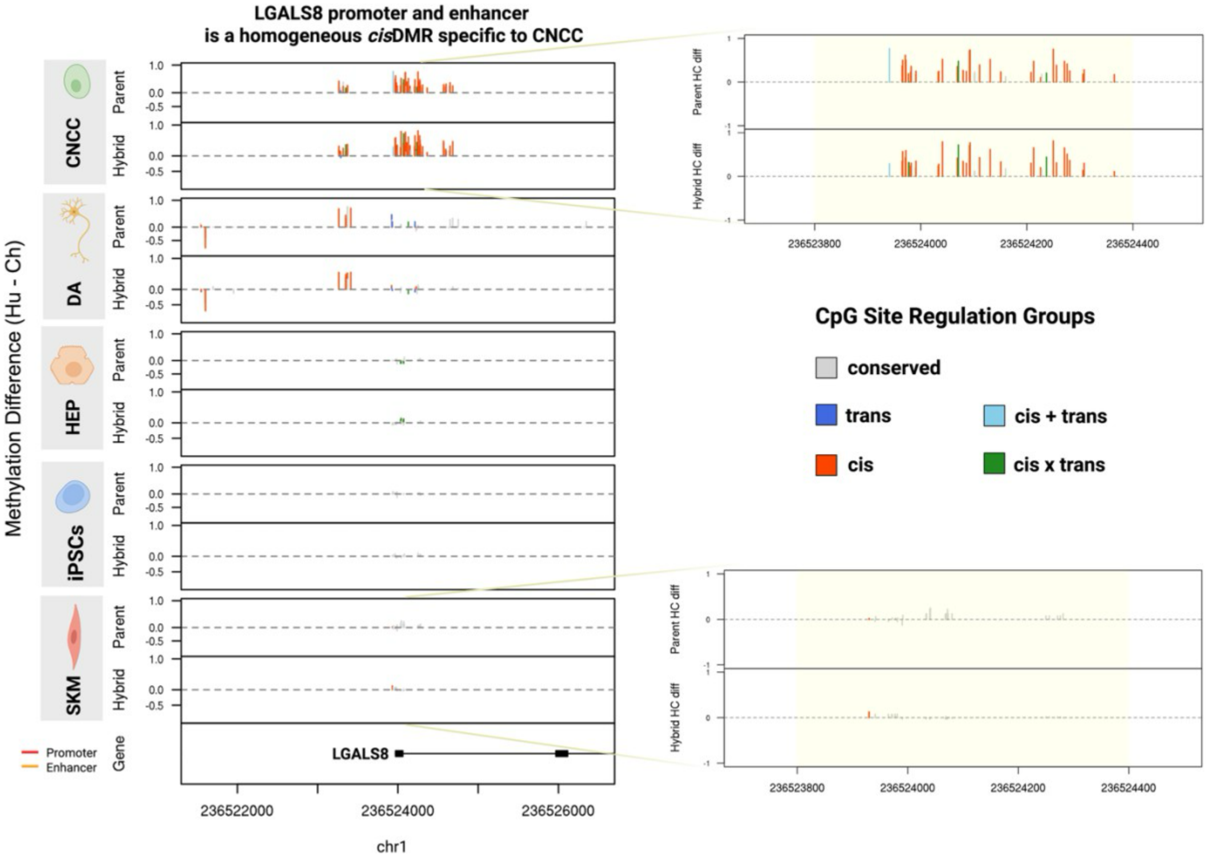


**Figure 4—figure supplement 2. *LGALS8* promoter and enhancer is a homogeneous *cis*-DMR specific to CNCC.** Differences of fractional methylation of human and chimpanzee in hybrid and parental cell types are compared. Each vertical line represents methylation difference of a single CpG site, colored by its regulation group. From Figure 4E, it is evident that the *LGALS8* promoter and enhancer region shows allele-specific methylation exclusively in CNCC while showing no allelic differences in other cell types. As seen from this figure, the allele-specific methylated region is also differentially methylated in CNCC parental cells and the majority of individual CpG sites are characterized as cis, meaning that this region is a *cis*-DMR, whereas other cell types show conserved methylation in the same region. See Supplemental File 3 for detailed statistics.

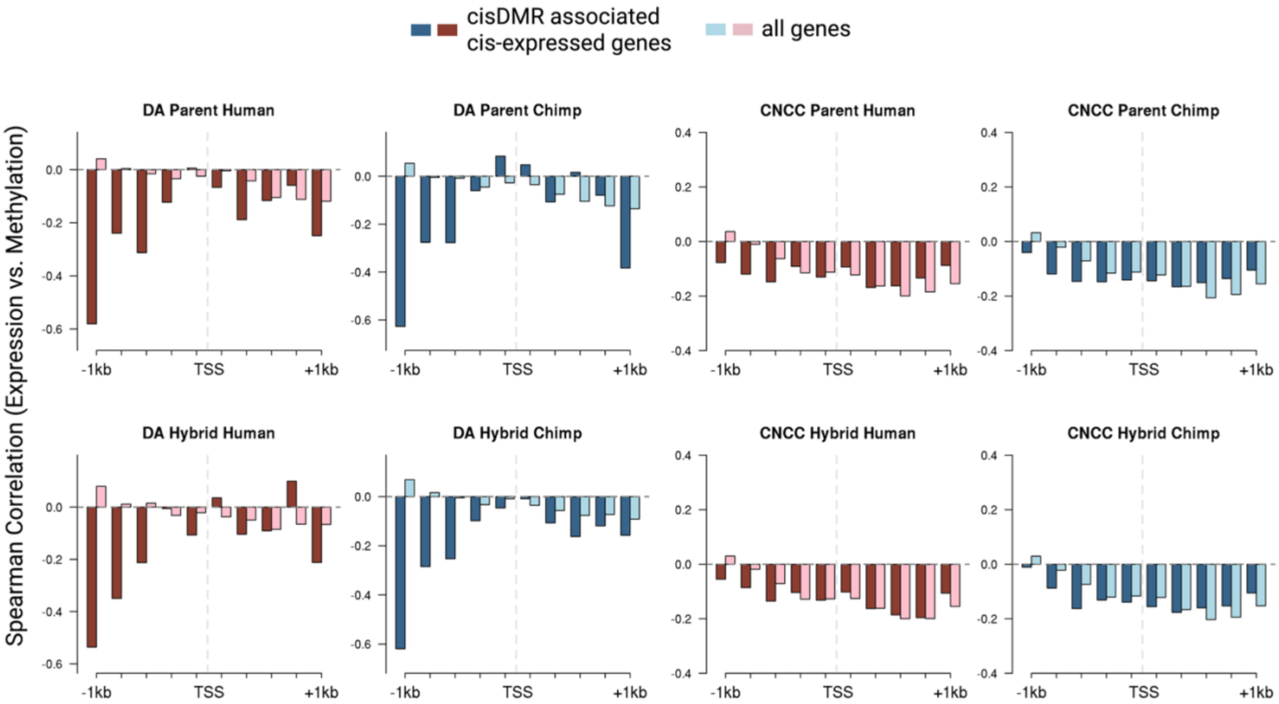


**Figure 4—figure supplement 3. Spearman correlation of expression and methylation for CNCC and DA.** Binned expression-methylation correlation profile of genes with both methylation and expression under cis-regulation (dark red and steel blue) and all genes regardless of regulation type (light pink and light blue). Two kb surrounding each transcription start site (TSS) is shown, split into 10 bins of 200 bp where individual CpG sites are pooled and averaged as fractional methylation values and correlated with expression values of the neighboring genes (in TPM). A weak but significant negative correlation is seen between expression and methylation levels near the promoter regions slightly upstream of TSS, consistent with past studies. When restricting to only the genes with cis-regulated methylation and cis-regulated expression, the correlation is stronger, especially in the bin 800 bp - 1 kb upstream of the TSS in DA cells. See Supplemental Files 5 and 6 for significance and detailed statistics.

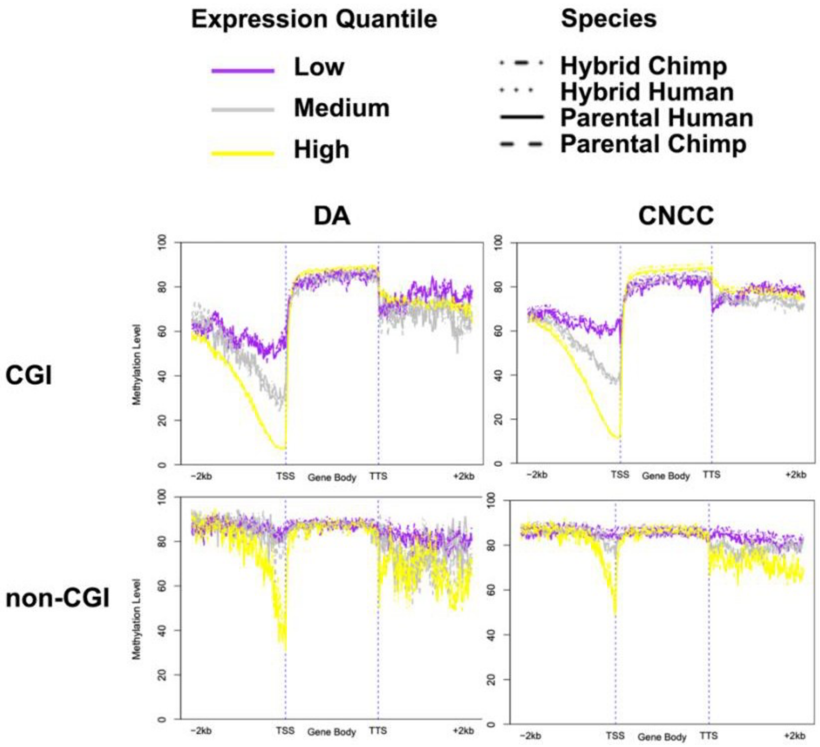


**Figure 4—figure supplement 4. CpG island (CGI) promoters show stronger repressive methylation-expression coupling than non-CGI promoters.** Genes were stratified into three equally sized groups based on their expression levels (TPM), and average methylation level of each group is shown in 100 bins across 2000bp upstream of transcription start site (TSS), gene body (all gene bodies scaled to be equal length in this plot), as well as 2000bp downstream of transcription termination site (TTS). A general pattern follows for CGI promoters that high expression is associated with low methylation, medium expression is associated with intermediate methylation levels, and low expression is associated with high methylation, whereas non-CGI promoters shows less stratification. Analysis of both CGI and non-CGI promoters revealed that genes are generally hypomethylated in promoter regions, with highly expressed genes showing a higher degree of hypomethylation in promoters, consistent with the established role of promoter methylation in transcriptional repression. However, CGI promoters showed more pronounced stratification, with highly expressed genes displaying significantly lower promoter methylation levels compared to lowly expressed genes.

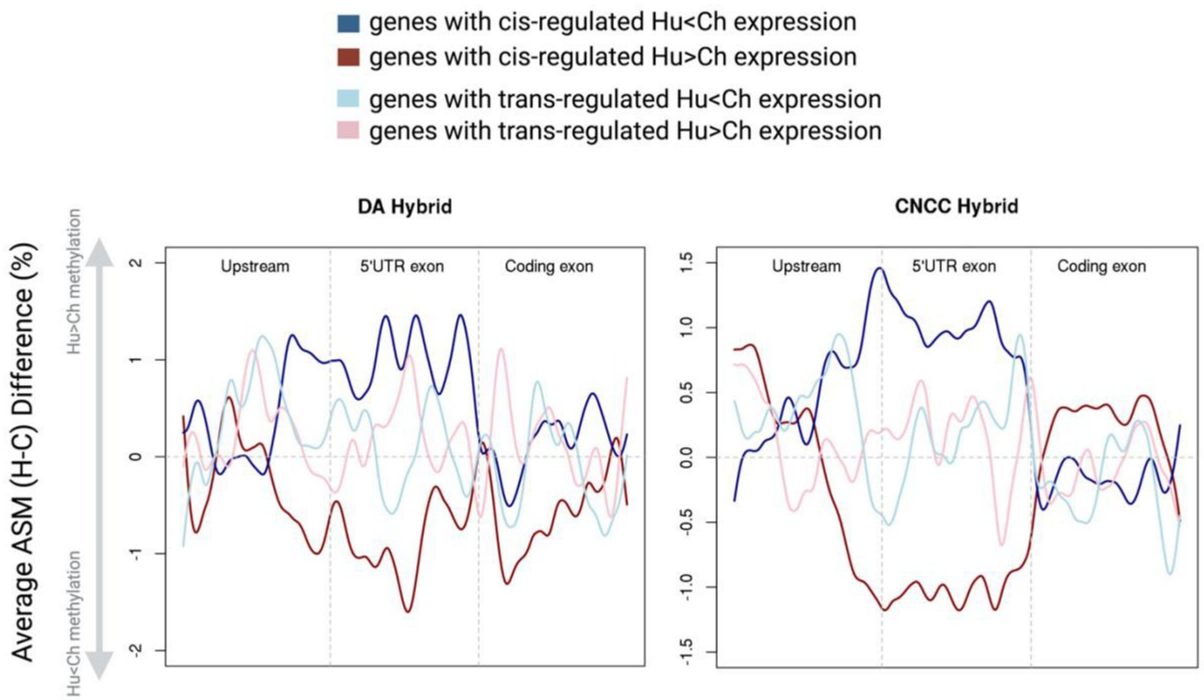


**Figure 4—figure supplement 5. Genes with cis-regulated expression are more likely to exhibit repressive patterns (i.e., higher methylation associated with lower expression) than those with trans-regulated expression**. The upstream regions (1000bp upstream of TSS), gene bodies and downstream regions (1000bp downstream of TTS) are each splitted into 100 equal bins where individual CpG sites are pooled and averaged as fractional methylation values. In genes with cis-regulated gene expression, the ones with Hu<Ch expression shows Hu>Ch methylation on average across ~1000bp upstream of and including 5' UTR region whereas genes with Hu>Ch expression shows Hu<Ch methylation. In genes with trans-regulated gene expression, the repressive pattern is not observed. This is consistent with DNA methylation acting as an upstream cis-regulatory mechanism.

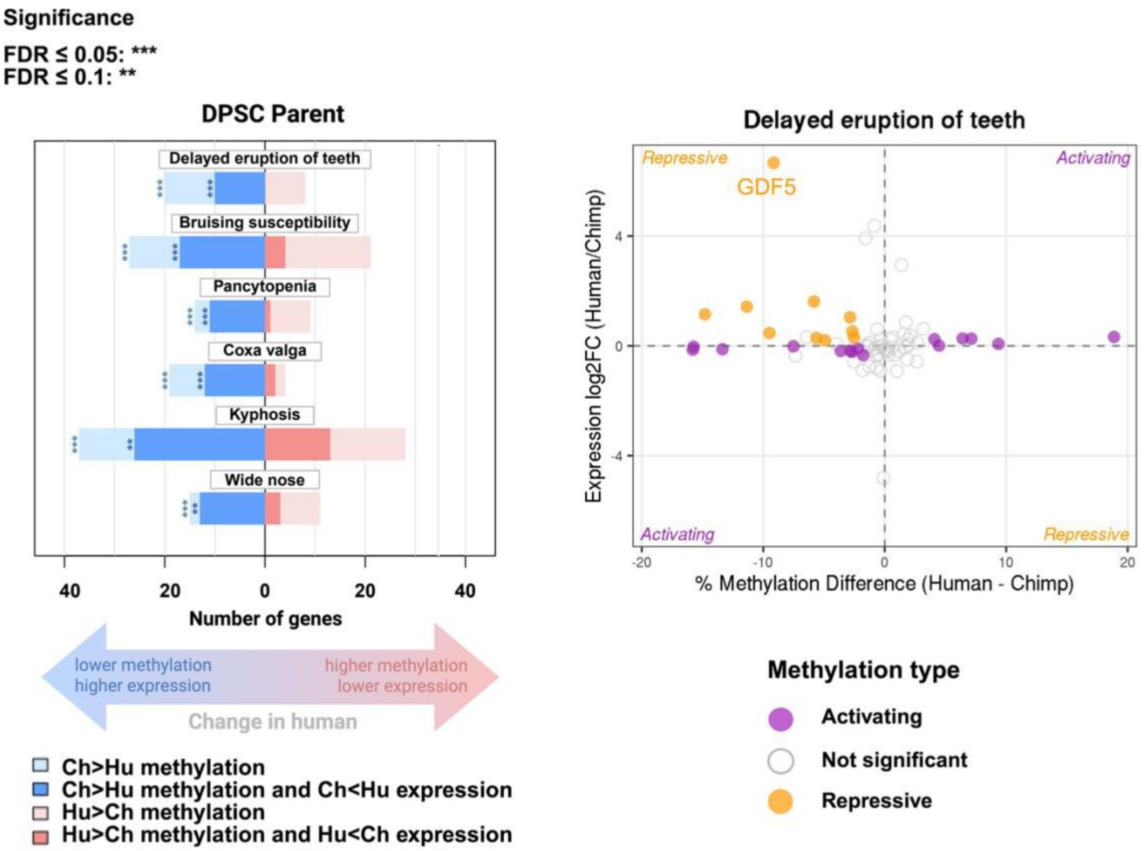


**Figure 5—figure supplement 1. Sign test on DPSC parental methylation and expression data shows significant bias for genes related to delayed eruption of teeth**. Left panel: The length of the bars indicates the number of genes in a category with allele-specific expression or methylation in each direction (human or chimpanzee-biased). The bars in bolded colors represent the number of genes within the pathway where higher methylation is associated with repressed gene expression, whereas lighter colors represent genes where higher methylation is associated with increased gene expression. Right panel: Differential expression log fold-changes are plotted against methylation. Notably, in pathway “delayed eruption of teeth”, *GDF5* has >30-fold higher expression and ~10% lower promoter methylation in human compared to chimpanzee DPSCs. Since the sign test assumes independence of gene expression changes within a pathway, results on parental data (which do not exclude trans-acting factors that may be shared across multiple genes) are suggestive but cannot reject the null hypothesis of neutral evolution. DPSC parental samples had a strong genome-wide human bias in promoter methylation, with 68% of promoters showing human-biased methylation compared to 32% chimp-biased, increasing the significance of gene sets with chimp-biased methylation. See Supplemental File 7 for detailed statistics.
